# Mechanical Loading due to Muscle Movement Regulates Establishment of the Collagen Network in the Developing Murine Skeleton

**DOI:** 10.1101/2022.06.23.497302

**Authors:** S. Ahmed, A.V. Rogers, N. C. Nowlan

## Abstract

Mechanical loading is critical for collagen network maintenance and remodelling in adult skeletal tissues, but the role of loading in collagen network formation during development is poorly understood. We test the hypothesis that mechanical loading is necessary for the onset and maturation of spatial localisation and structure of collagens in prenatal cartilage and bone, using *in vivo* and *in vitro* mouse models of altered loading. The majority of collagens studied were aberrant in structure or localisation, or both, when skeletal muscle was absent *in vivo*. Using *in vitro* bioreactor culture system, we demonstrate that mechanical loading directly modulates the spatial localisation and structure of collagens II and X. Furthermore, we show that mechanical loading *in vitro* rescues aspects of collagens II and X development from the effects of fetal immobility. In conclusion, our findings show that mechanical loading is a critical determinant of collagen network establishment during prenatal skeletal development.

**Teaser (One line):** Mechanical loading is required for normal establishment and maturation of key collagens during prenatal skeletal development.

## Introduction

The matrix of articular cartilage has an extraordinarily complex structure and composition, which enables it to sustain high loads and facilitate almost frictionless movements throughout a lifetime, while the extracellular matrix of bone facilitates the strength and toughness of the tissue. The tensile properties of cartilage and bone depend on an intricate collagen network comprising multiple types of individual collagens. Numerous syndromes and disorders in which specific collagens are abnormal impact on skeletal development (*1*), including osteogenesis imperfecta (*2*) and chondrodysplasias (*3*). Changes in the collagen network also play a role in age-related degeneration and the aetiology of osteochondral disorders, including osteoarthritis (*4*).

Articular cartilage and bone tissues arise from a cartilage template which forms in the early embryo (*5*). As prenatal development of a long bone progresses, most of the cartilage is replaced through endochondral ossification. While articular cartilage was previously believed to form appositionally from the cartilage template (*6, 7*), recent work has demonstrated that articular cartilage develops mainly interstitially (*8*). Therefore, while the matrix of the skeletal tissues is substantially remodelled over ontogeny, the origins of the mature articular cartilage and bone matrix lie in prenatal development. In prior work, we revealed the dynamic changes in collagen fibre organisation and complexity that occur over prenatal skeletal development, particularly in collagens II, X and XI (*9*). The dynamism of collagen network emergence is likely to be critical to the functional and biological properties of the mature skeletal tissues, but the mechanisms underlying the increasing complexity of the multiscale collagen network over development are unknown.

Mechanical loading plays a critical role in the maintenance and health of mature and ageing skeletal tissues (*10*). Mechanical loading is also critical to prenatal and postnatal skeletal development. When fetal movements are reduced, restricted, or (in extreme cases) absent, skeletal malformations can result including joint shape abnormalities (e.g., developmental dysplasia of the hip and arthrogryposis), congenital scoliosis, and thin, hypo-mineralised bones [reviewed in (*11*)]. While no prior studies have described the effects of abnormal fetal movements on the matrix of skeletal tissues, there is evidence from studies on postnatal development that mechanical loading is important for emergence of the collagen network. It has been postulated that mechanical loading due to physical activity regulates the emergence of complexity in the collagen network of postnatal articular cartilage (*5, 12, 13*). For example, collagen content and organisation differ between differentially loaded sites of the equine joint cartilage (*5*), and exercise regimes have been shown to influence collagen organisation in the articular cartilage of adolescent animals (*5, 14*). Mechanical loading also influences the collagen network at the cellular level; tensile forces between collagen fibre bundles and the cell membrane protrusions (“fibripositors”) from which collagen fibrils exit the cell have been proposed as drivers of collagen fibril formation, orientation, and organisation (*15*). In this study, we hypothesise that mechanical loading is a critical factor in the initiation and maturation of the collagen network during prenatal skeletogenesis. To test this hypothesis, we first studied the effects of skeletal muscle contractions on the localisation and structure of key skeletal collagens during prenatal development, using the Splotch-delayed mouse model which has no skeletal muscles (Pax3^spd-/-^, or “muscleless-limb”). Next, we determined if the effects of absent muscle loading on the collagen network are mediated directly by changes in mechanical loading, rather than by changes in the biochemical environment when skeletal muscle is missing. Fetal limb explants were cultured under static or dynamically loaded conditions in a mechano-stimulation bioreactor, to assess the impact of mechanical loading on collagen network development independently of muscle contractions. Finally, we tested if *in vitro* application of mechanical loading could “rescue” the effects of fetal immobility on collagen network development. Understanding the contribution of mechanical loading to collagen network establishment and maturation provides insight into skeletal development and disease, and has the potential to lead to therapies for babies and children affected by reduced or abnormal movements.

## Results

### Collagens I, II, V, VI, X and XI are abnormal in the developing rudiment when muscle is absent

In order to investigate the role of skeletal muscle contractions *in utero* on prenatal skeletal development, we used a mouse line in which no limb skeletal muscles form in embryos homozygous for the mutation; Pax3^spd-/-^, or Splotch-delayed, referred to herein as “muscleless-limb” mice. Three significant developmental stages of rudiment development were studied; Theiler Stage (TS)22 (*16*) (typically embryonic day (e)13.5), the latest stage at which the rudiment is fully cartilaginous, TS25 (typically e16.5) when the growth plate and mineralised regions are present, and TS27 (typically e18.5) the latest prenatal stage that can be reliably analysed. Muscleless-limb embryos and their normal littermates were harvested and staged. Seven collagen types (collagens I–III, V, VI, X and XI) were studied with immunofluorescence on cryosections of the humerus and imaged with confocal microscopy. The selected seven collagens were the same as those we characterised for normal development, except for collagen IX, which was not included in the current study due to the lack of any observed changes over normal development (*9*). Confocal images were processed using FIJI (*17*) and qualitatively analysed to identify differences in the spatial localisation between wild type and muscleless-limbs over the developmental time course.

The absence of skeletal muscle did not lead to any obvious changes in collagen III at any of the stages studied (Supplementary Figure 3). All other collagens studied were affected by the lack of skeletal muscle in terms of their structural organisations and/or their spatial localisation, as described in detail below. Results shown are representative of all repeats unless otherwise described, and the complete dataset is provided in Supplementary Figures 1–8.

**Collagen I** is normally localised to the perichondrium and the mineralising cartilage and plays an important role in bone mass and strength (*18*). In the TS22 wild types, collagen I localisation was concentrated in the perichondrium (Figure 1A, a), and at the proximal epiphysis (Figure 1A, arrow). In the muscleless-limbs at TS22, there was localisation of collagen I throughout much of the diaphysis, lacking specificity to the perichondrium (Figure 1B, b). At TS25 and TS27, collagen I localisation in the wild types was concentrated in the mineralising region and in the perichondrium (Figure 1C, F). One of the two muscleless-limbs studied at TS25 (“Muscleless 1”) exhibited diffuse collagen localisation throughout the cartilaginous region (Figure 1D), and did not exhibit a clear cartilage to mineralisation interface (Figure 1D, dashed line) or the distinct collagen I fibre organisation (Figure 1d) seen in the wild type rudiments (Figure 1C, c). Collagen I localisation and organisation in the other TS25 muscleless-limb (“Muscleless 2”, Figure 1E, e) were similar to the TS25 wild types (Figure 1C, c). At TS27, both structure and localisation of collagen I were abnormal when skeletal muscles were absent. TS27 muscleless-limbs lacked collagen I localisation in the distal perichondrium (compare arrows in Figure 1F & Figure 1G). Punctate localisation of collagen I was seen throughout the non-mineralised cartilage in the TS27 muscleless-limbs (Figure 1g) but not in the wild types (Figure 1f). In the mineralising region, the collagen I fibres in the muscleless-limbs appeared finer and less connected than those of wild types (Figure 1h, i). Therefore, when skeletal muscles were lacking, the structure and localisation of collagen I was aberrant, with the consequences changing with developmental stage.

**Figure 1:**
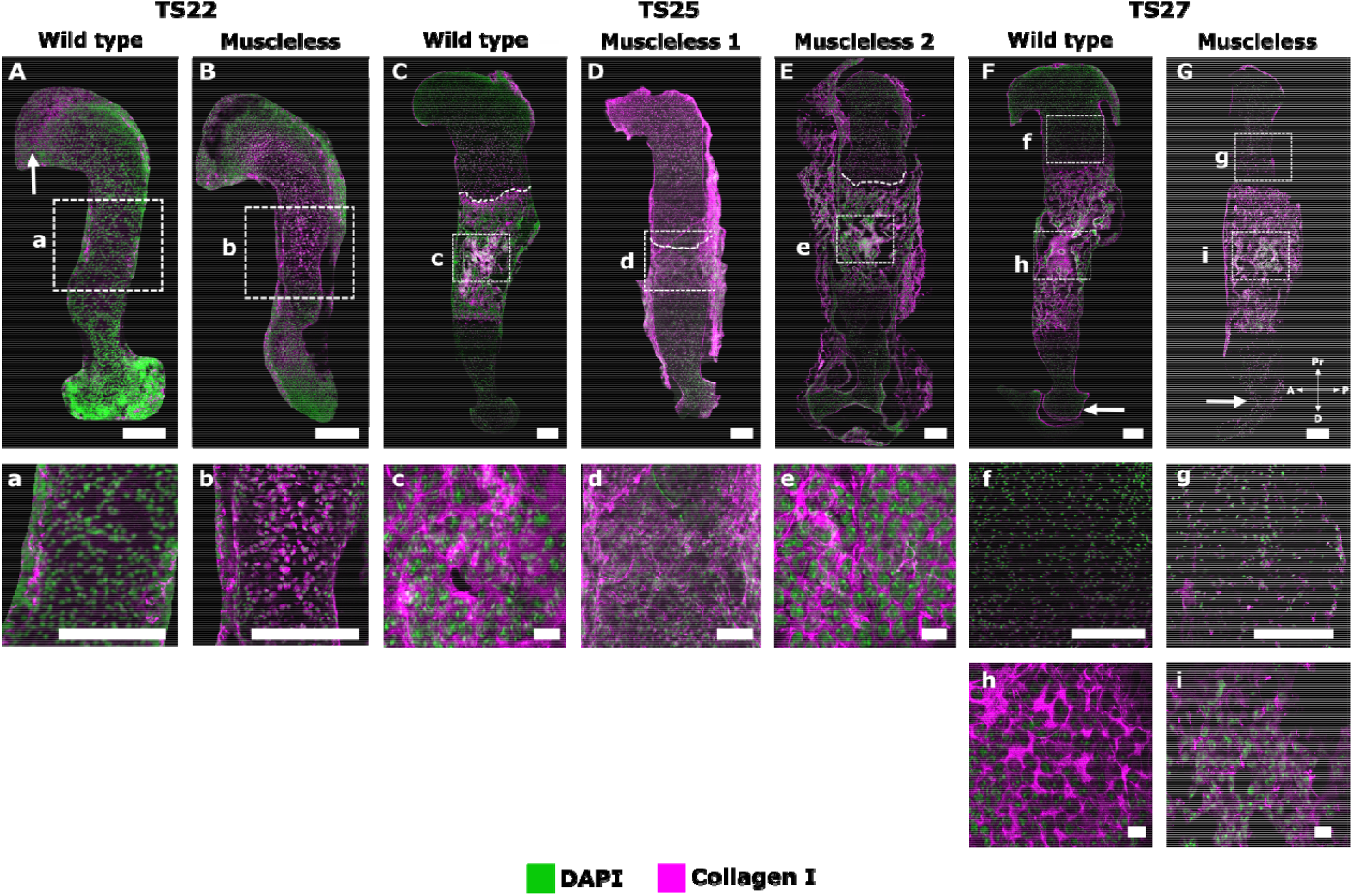
Both structure and localisation of collagen I were abnormal in muscleless-limbs, with the effects varying according to the developmental stage. A–B, a–b: In the TS22 wild types, collagen I localisation was concentrated in the perichondrium (A, a), and at the proximal epiphysis (A, arrow). In the muscleless-limbs at TS22 (B, b), there was localisation of collagen I throughout much of the diaphysis, lacking specificity to the perichondrium. C–E, c–e: At TS25, collagen I localisation in the wild types was concentrated in the mineralising region and in the perichondrium (C). One of the muscleless-limbs at TS25 (“Muscleless 1”) had abnormal collagen localisation throughout the cartilaginous region (D), lacked a clear cartilage to mineralisation interface (D, dashed line), and also lacked the distinct collagen I fibre organisation (d) seen in the wild type rudiments (C, c). The other TS25 sample “Muscleless 2” (E, e) had similar collagen I localisation and organisation to the wild type. Dashed lines in C–E demarcate mineralising and non-mineralised regions. F–G, f–g: At TS27, both structure and localisation of collagen I were abnormal when skeletal muscles were absent. Muscleless-limbs lacked collagen I localisation in the distal perichondrium (compare arrows in F & G). Punctate localisation of collagen I was seen throughout the non-mineralised cartilage in the muscleless-limbs (g) but not in the wild types (f). In the mineralising region, collagen I fibres in the muscleless-limbs were finer and less interconnected than those of wild types (h, i). Except for the scale bars in (c), (e), (h), and (i), which show 10 μm, all scale bars are 200 μm. Sample numbers: TS22 wild type/muscleless (n=3), TS25 wild type (n=3), TS25 muscleless (n=2), TS27 wild type (n=4), TS27 muscleless (n=3).

**Collagen II** is the major collagen of cartilage, normally present in the non-mineralized regions of cartilaginous rudiments and, later in prenatal development, the perichondrium (*9*). Collagen II is necessary for the proper formation of articular cartilage, the epiphyseal growth plate, endochondral bone and the intervertebral disc (*19, 20*). Both spatial distribution and structural organisation of collagen II were affected by the lack of skeletal muscles. At TS22, the muscleless-limbs exhibited widespread perichondral localisation (including the future articular cartilage region), whereas wild type limbs had only mild collagen II staining in the perichondrium at this stage (Figure 2A, B arrows). A meshwork-like pattern was already evident in the muscleless-limbs, while no such pattern was obvious in the wild types (Figure 2a, b). At TS25, perichondral localisation of collagen II in the muscleless-limbs was much more pronounced and intense than in the wild type limbs (Figure 2C & D, arrows). TS25 muscleless-limbs exhibited a prominent collagen II meshwork in the proximal humerus with thick interconnected bundles (Figure 2d), which was not observed in the wild types (Figure 2c). The hypertrophic regions of the muscleless rudiments had thick bundles of collagen II localisation in the longitudinal septa (Figure 2f, star) with thinner bundles in the transverse septa (Figure 2f, arrowhead), a pattern that was much more pronounced compared to that seen in the wild types (Figure 2e, star and arrowhead). In the TS25 mineralised regions, muscleless-limbs had a fibrous collagen II organisation (Figure 2h) whereas no collagen II was present in the mineralised regions in the wild type rudiments (Figure 2g). By TS27, no differences in collagen II localisation between muscleless-limbs and wild types were evident (Figure 2E, F). However, the characteristic meshwork pattern of collagen II in the proximal humerus was almost completely lost in the muscleless-limbs, but fully established in the wild types (Figure 2i, j). A distinctive septal organisation was established throughout the growth plates of the TS27 wild types (Figure 2k) while no longer present in the muscleless-limbs (Figure 2l). Mild collagen II immunopositivity was observed in the mineralised region of the muscleless-limbs (Figure 2n) but not in the wild types (Figure 2m). Therefore, early in development, the absence of skeletal muscle led to abnormal collagen II localisation in the perichondrium, and a precocious structural arrangement in the epiphyseal cartilage. However, by TS27, the muscleless-limbs had lost all of the structural arrangement of collagen II, the same stage at which the wild type limbs have a pronounced structure.

**Figure 2:**
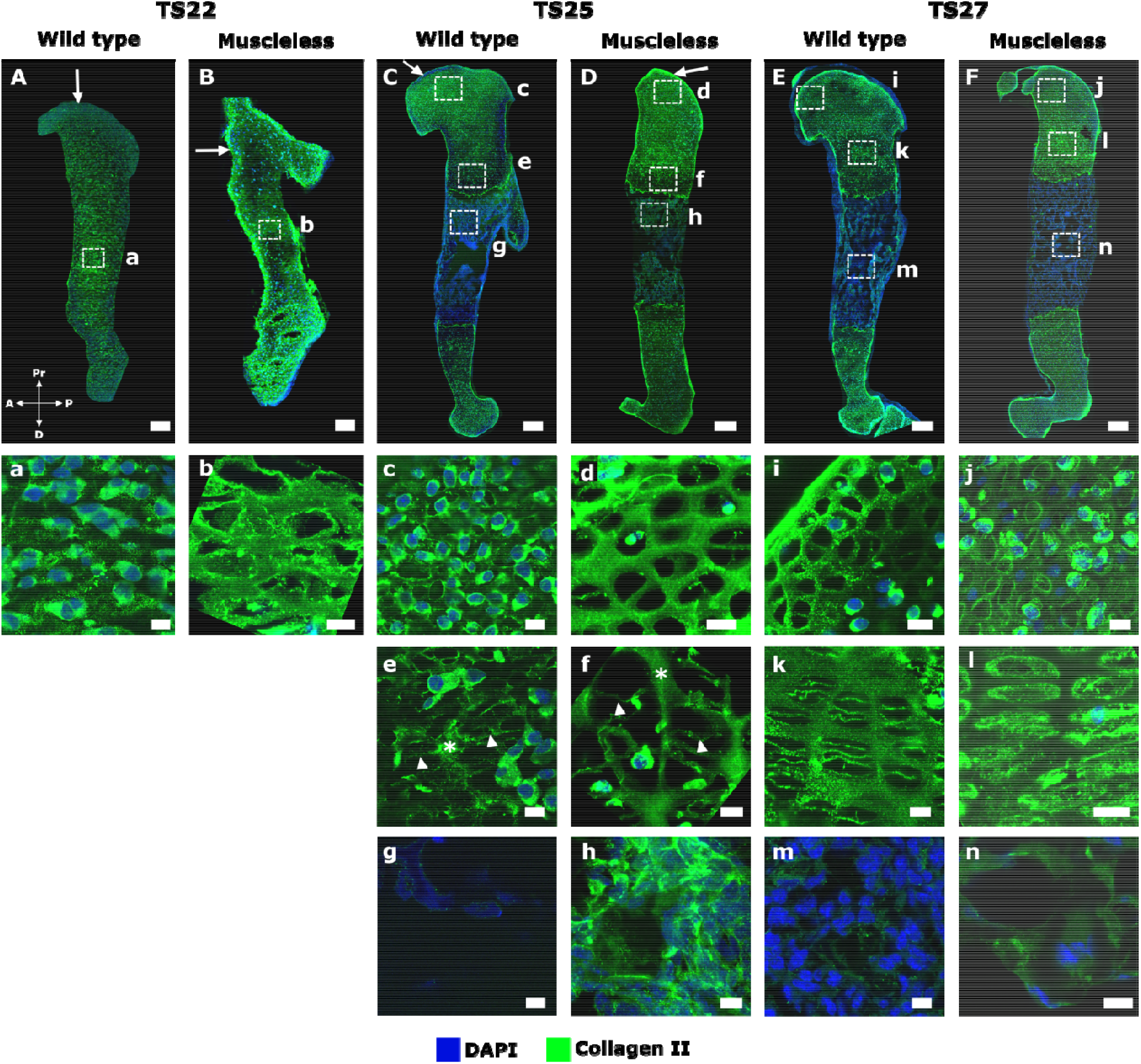
Collagen II exhibited an initially more structured organisation in muscleless -limbs than in wild type limbs, but this structure was lost by TS27, when the wild type characteristic meshwork pattern was fully established. A–B, a–b: At TS22, muscleless-limbs exhibited widespread perichondral localisation (including the future articular cartilage region), whereas wild type limbs had only mild collagen II staining in the perichondrium at this stage (arrows A, B). A meshwork-like pattern was already evident in the mid-diaphysis of muscleless-limbs, while no such pattern was obvious in the wild types (a, b). C–D, c–h: At TS25, perichondral localisation of collagen II in the muscleless-limbs was more pronounced than in the wild type limbs (arrows in C, D). Muscleless-limbs exhibited a prominent collagen II meshwork in the proximal humerus with thick interconnected bundles (d), not observed in the wild types (c). The hypertrophic regions of the muscleless-limbs had thick bundles of collagen II localisation in the longitudinal septa (f, star) with thinner bundles in the transverse septa (f, arrowhead), a pattern that was more pronounced compared to that seen in the wild types (e). In the TS25 mineralising regions, muscleless-limbs had a fibrous collagen II organisation (h), whereas no collagen II was present in the mineralising regions in the wild type rudiments (g). E–F, i–n: By TS27, no differences in collagen II localisation between muscleless-limbs and wild types were evident (E, F). The characteristic meshwork pattern of collagen II in the proximal humerus was almost completely lost in the muscleless-limbs, but fully established in the wild types (i, j). A distinctive septal organisation was established throughout the growth plates of the TS27 wild types (k) while no longer present in the muscleless-limbs (l). Mild collagen II immunopositivity was observed in the mineralised region of the muscleless-limbs (n) but not in the wild types (m).). Scale bars 88 µm (A–B), 200 µm (C–G) and 10 µm (a–n). Sample numbers: TS22 wild type (n=4), TS22 muscleless (n=3), TS25 wild type (n=3), TS25 muscleless (n=2), TS27 wild type/muscleless (n=3).

**Collagen V** is important for the regulation of collagen I fibre number and diameter (*21-24*) and therefore plays an important role in bone development. Several aspects of collagen V localisation were abnormal in the muscleless-limbs, while no pronounced abnormalities in structure were evident at any stage (Figure 3). At TS22, collagen V was present throughout the muscleless-limbs (Figure 3B), whereas there was consistent diaphyseal localisation in the wild type rudiments, without any epiphyseal localisation (Figure 3A). At TS25, there were no pronounced abnormalities in localisation of collagen V (Figure 3C, D). However, at TS27, the muscleless-limbs had more pronounced collagen V localisation in the perichondrium and the proximal joint region (Figure 3F, arrowheads) compared to the wild types (Figure 3E, arrowheads). Therefore, the absence of muscles led to abnormal localisation of collagen V at two of the three stages examined, with no obvious effects on structure.

**Figure 3:**
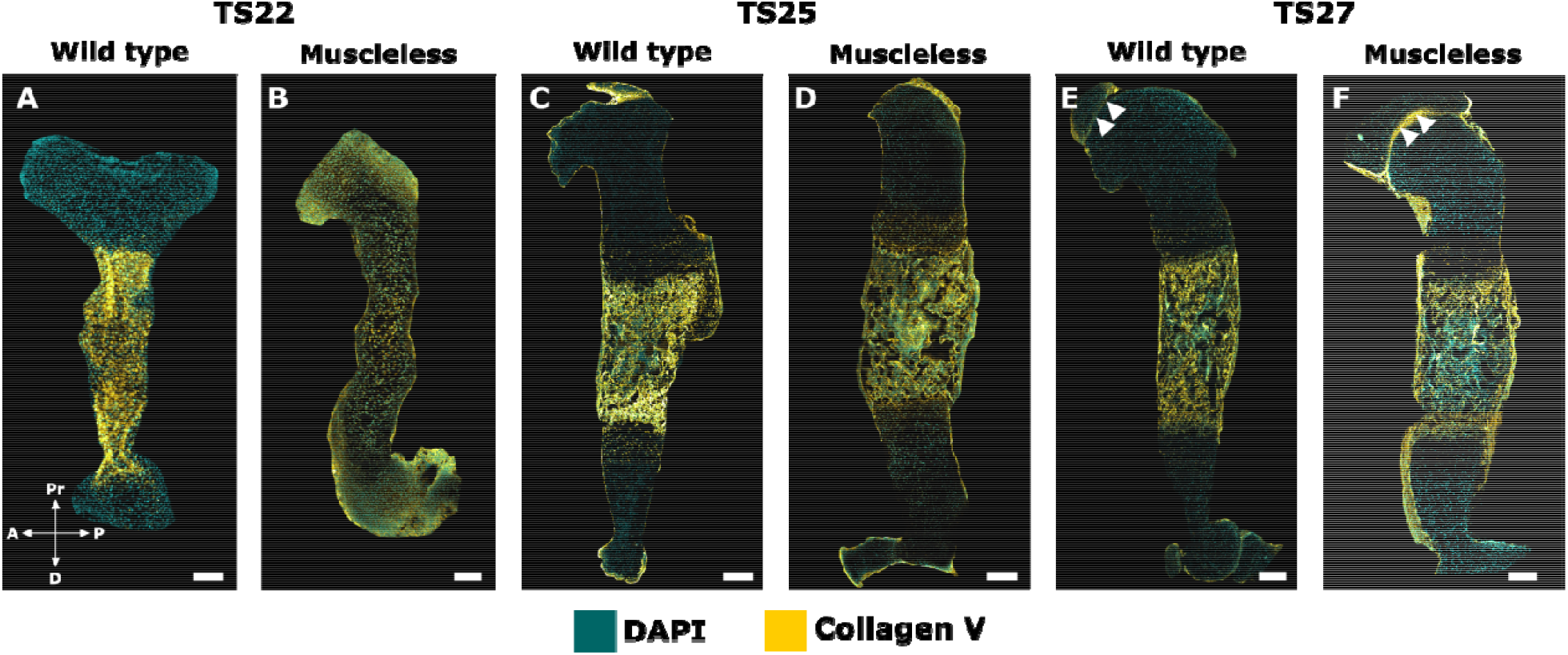
Collagen V localisation was aberrant at TS22 and TS27 in muscleless-limbs, with no evident structural abnormalities. A, B: At TS22, collagen V was present throughout the muscleless-limbs (B), whereas there was consistent diaphyseal localisation in the wild type rudiments, without any epiphyseal localisation (A). C, D: At TS25, there were no pronounced abnormalities in localisation of collagen V. E, F: At TS27, the muscleless-limbs had more pronounced collagen V localisation in the perichondrium and the proximal joint region compared to the wild types (arrowheads in E & F). Scale bars 200 μm. Sample numbers: TS22 wild type/muscleless (n=3), TS25 wild type (n=3), TS25 muscleless (n=2), TS27 wild type/muscleless (n=3).

**Collagen VI** is localised to the pericellular matrix and delineates chondron structure (*25*). The pericellular matrix is a key player in chondrocyte mechanotransduction (*26*). While the overall spatial distribution of collagen VI was not visibly affected by the lack of skeletal muscle at TS22 and TS25 (Figure 4A–D), there was a pronounced increase in staining intensity of collagen VI in the proximal humerus in the muscleless-limbs at TS27 compared to the tight joint line localisation in the wild types at the same stage (Figure 4a, b, arrows). Chondron shapes appeared normal at TS22 and TS25 (Supplementary Figure 5), but structural differences in chondron shape were observed in the proximal humerus by TS27. While the TS27 chondrons had an ovoid convexity in the wild types (Figure 4c), the muscleless-limb chondrons at the same stage were more cylindrical in shape (Figure 4d). Therefore, a lack of skeletal muscle affected collagen VI only at latest stage examined, with increased joint line localisation and altered chondron shape.

**Figure 4:**
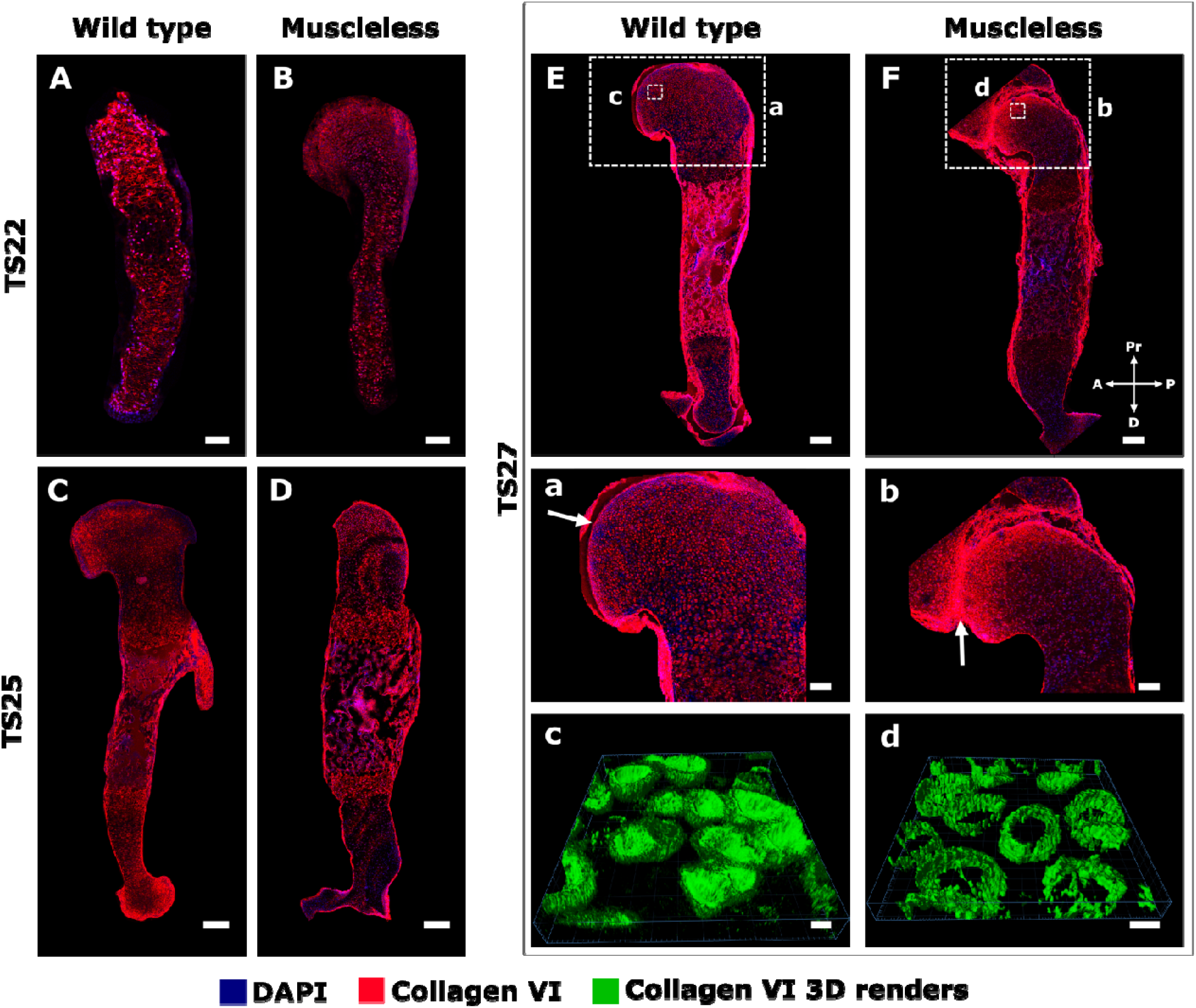
Effects of absent skeletal muscle on collagen VI were only detected at the last stage studied, with increased joint line localisation and altered chondron morphology in the TS27 muscleless-limbs. A–D: The overall spatial distribution of collagen VI was not visibly affected by the lack of skeletal muscle at TS22 (A, B) or at TS25 (C, D). E, F, a–d: At TS27, there was a pronounced increase in staining intensity of collagen VI in the proximal humerus in the muscleless-limbs compared to the tight joint line localisation in the wild types at the same stage (a, b, arrows). While the TS27 chondrons had an ovoid convexity in the wild types (c), the muscleless-limb chondrons at the same stage were more cylindrical in shape (d). Scale bars 200 μm (A–F), 100 µm (a– b) and 5 µm (c–d). Sample numbers: TS22 wild type/muscleless (n=3), TS25 wild type (n=3), TS25 muscleless (n =2), TS27 wild type/muscleless (n=3).

**Collagen X** is an important component of the growth plate (*27*) and, during development, is normally present in the pre-hypertrophic and hypertrophic growth plate regions (*9*). Both localisation and structure of collagen X were affected by the absence of skeletal muscle. In wild type limbs, collagen X was strongly localised to the mineralising cartilage, with some sparse punctate localisation in the non-mineralised cartilage (Figure 5A, C, E, c & g). Punctate localisation in the non-mineralised cartilage was much more widespread and pronounced in the muscleless-limbs at TS25 and TS27 than in the wild types (compare Figure 5 d to c, and h to g). In the growth plates of the wild type limbs, collagen X staining was mainly extracellular (Figure 5a, e & i), while in the muscleless-limbs, there was a tendency across all three stages for increased punctate, intracellular localisation in the growth plates, as indicated with arrows in Figure 5b, f, & j. The structure of collagen X was also affected by the lack of skeletal muscle. At TS22, both wild type and muscleless-limbs had a hexagonal collagen X structure in the mid-diaphysis (Figure 5a, b). However, the wild type collagen X structure at TS22 had an intricate weave with an obvious anterior-posterior grid-like pattern (Figure 5a), whereas the structure was more randomly oriented in the muscleless-limbs at the same stage (Figure 5b). In the wild types at TS25, collagen X exhibited a complex interlocking arrangement in the growth plate (Figure 5e), while once again the organisation in the muscleless-limbs was less structured (Figure 5f). In the TS27 wild type limbs, collagen X had a columnar arrangement in the pre-hypertrophic region, and a teardrop, or oblique, orientation in the hypertrophic regions (Figure 5i). In contrast, in the muscleless-limb rudiments at TS27, there was no complexity of structure within the different parts of the growth plate, with only an approximate columnar arrangement (Figure 5j). Therefore, the absence of skeletal muscle led to increased punctate localisation of collagen X in the non-mineralised cartilage, increased intracellular localisation in the mineralising cartilage, and pronounced effects on the structure of collagen X in the growth plate at all three stages examined.

**Figure 5:**
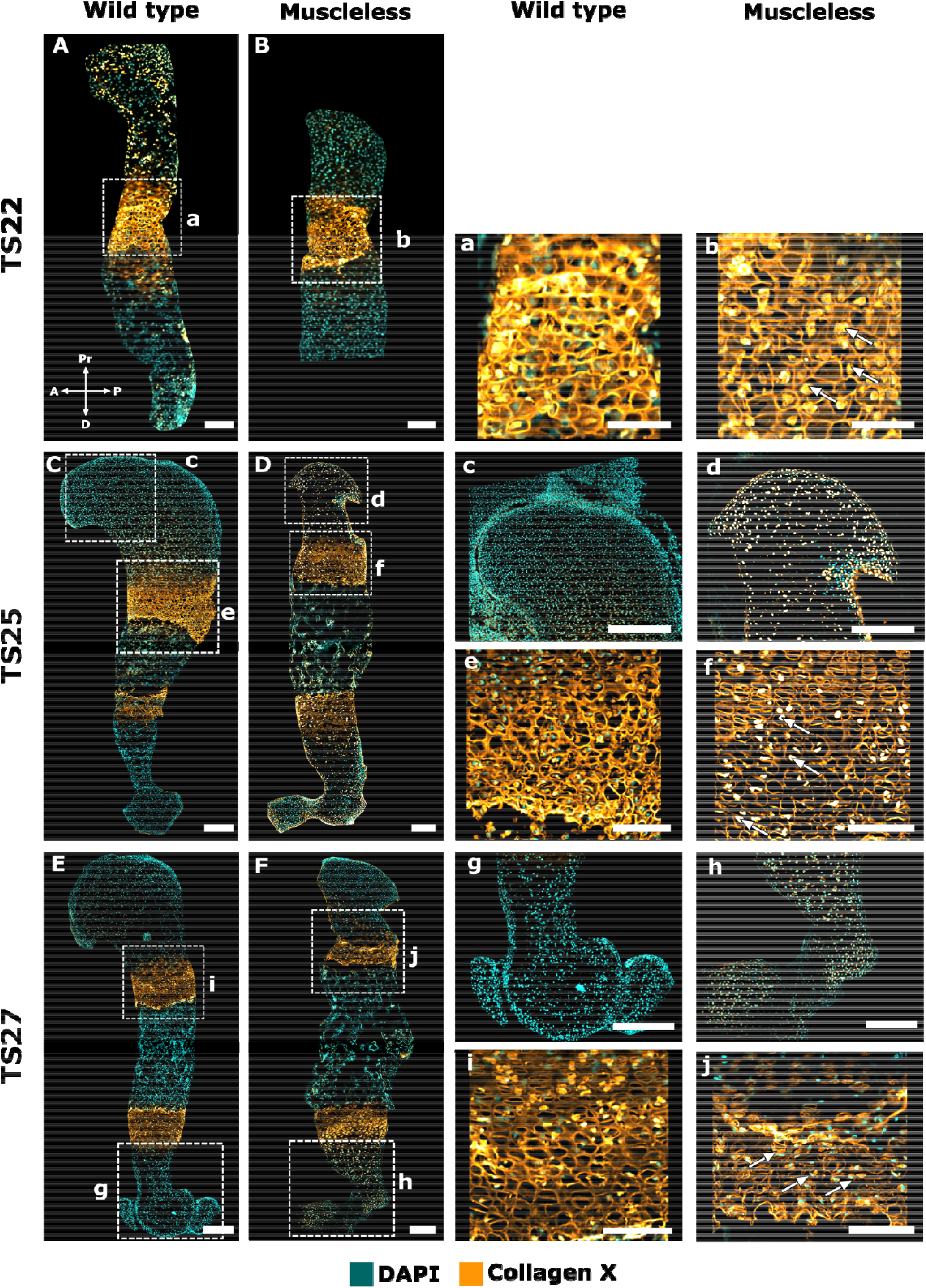
At all three stages studied, muscleless-limbs had increased intracellular localisation of collagen X, and aberrant growth plate structure. A, B, a, b: The wild type collagen X structure at TS22 had an intricate weave with an obvious anterior-posterior grid-like pattern (a), whereas the structure was more randomly oriented in the muscleless-limbs (b). The muscleless-limbs had increased punctate, intracellular localisation in the growth plates (b, arrows). C, D, c–f: At TS25, collagen X was strongly localised to the mineralising cartilage in wild types (C, c), while punctate localisation in the non-mineralised cartilage was widespread in the muscleless-limbs (D, d). As at TS22, the TS25 muscleless-limbs had increased punctate, intracellular localisation in the growth plates (f, arrows). There was a less defined interlocking arrangement of collagen X in the muscleless-limb growth plate (f) than in the wild type growth plate (e). E, F, g–j: At TS27, punctate localisation in the non-mineralised cartilage was still more pronounced and widespread in the muscleless-limbs (F, h) than in the wild types (E, g). TS27 muscleless-limbs had increased intracellular localisation in the growth plates (j, arrows). In the wild type limbs, collagen X had a columnar arrangement in the pre-hypertrophic region, and an oblique, orientation in the hypertrophic regions (i). In the muscleless-limbs, there was no complexity of structure within the different parts of the growth plate, with only an approximate columnar arrangement (j).Scale bars 200 μm (A–F), 65 µm (a–b), 170 µm (c) and 100 μm (d–j). Sample numbers: TS22 wild type (n=3), TS22 muscleless (n=2), TS25 wild type (n=3), TS25 muscleless (n =2), TS27 wild type/muscleless (n=3).

**Collagen XI** is important for chondrogenesis and has a protective role in maintaining healthy cartilage (*28*). It co-polymerises with collagen II (*29*) and, during development, is predominantly present in non-mineralized cartilage (*9*). No substantial differences in collagen XI structure or localisation were observed between wild type and the muscleless-limbs at TS22 (Figure 6A, B, a & b) or TS25 (Figure 6 C, D, c & d). However, by TS27, dramatic differences were observed between the wild type and the muscleless-limbs in all regions. In two out of the three samples, muscleless-limbs lacked the meshwork pattern in the proximal humerus and also lacked a distinct septal organisation in the growth plate regions (Figure 6E, F, e, f, h & i). In the third muscleless-limb sample, there was a complete absence of any collagen XI extracellular matrix arrangement (Figure 6G, g & j). Also at TS27, muscleless-limb rudiments had increased intracellular collagen localisation compared to wild types (Figure 6f, g, I & j arrows). Therefore, the structure and localisation of collagen XI initiated apparently normally when skeletal muscle was absent, but the structure of collagen XI was dramatically affected by the lack of skeletal muscle by TS27, as was the balance between extracellular and intracellular localisation.

**Figure 6:**
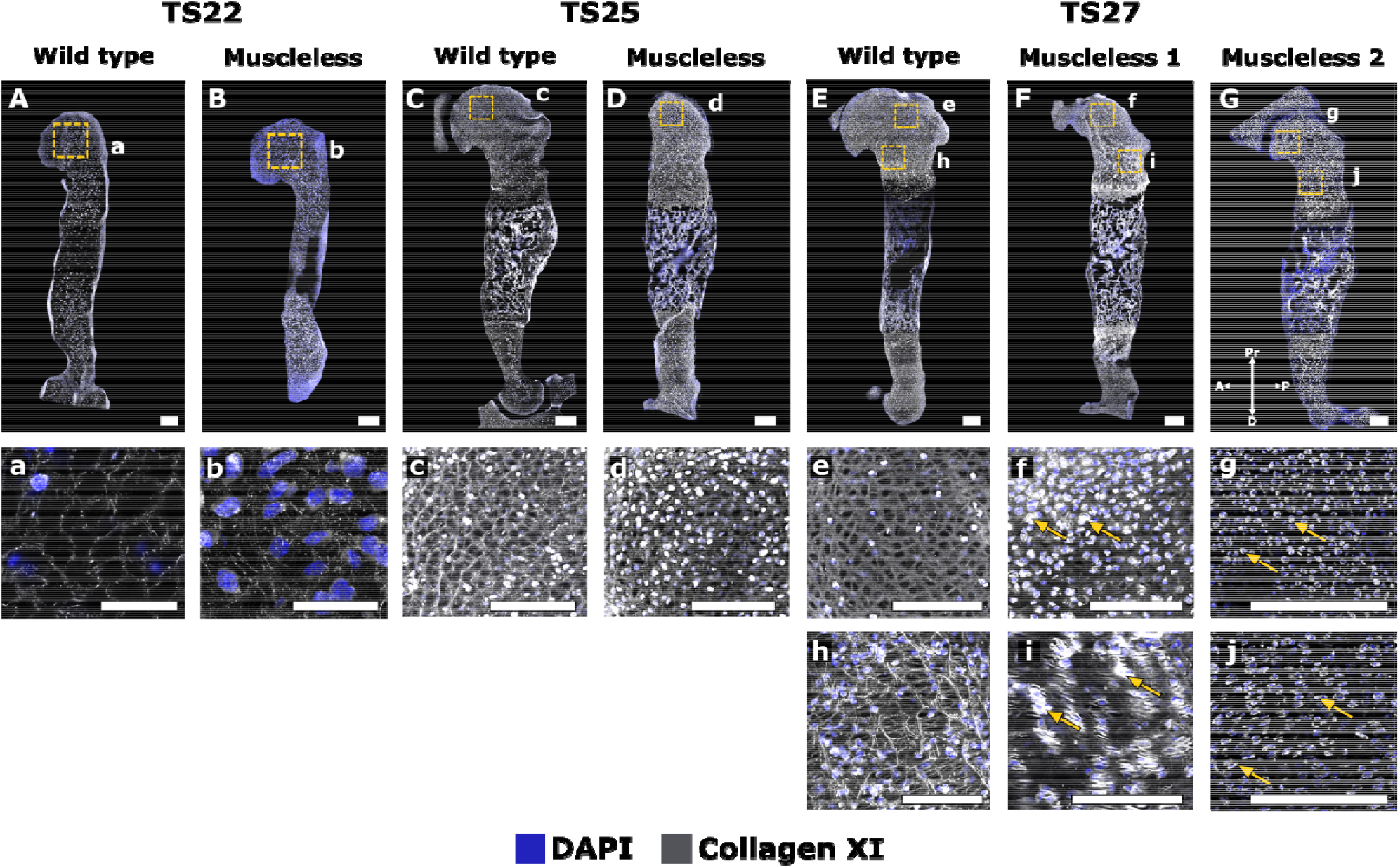
Effects of absent skeletal muscle on collagen XI were only detected at the last stage studied, with substantially altered structure in the TS27 muscleless-limbs. A–D, a–d: No substantial differences in collagen XI structure or localisation were observed between wild type and the muscleless-limbs at TS22 (Aa, Bb) or TS25 (Cc, Dd). E–G, e–j: At TS27, dramatic differences in collagen XI were observed between the wild type and the muscleless-limbs in all regions. In two out of the three samples, muscleless-limbs had no meshwork pattern in the proximal humerus and no distinct septal organisation in the growth plate regions (E, F, h & i). In the third muscleless-limb sample, there was a complete absence of any collagen XI extracellular matrix arrangement (G, g & j). TS27 muscleless-limb rudiments also had increased intracellular collagen localisation compared to wild types (arrows, f, g, i & j). Scale bars 200 μm. Sample numbers: TS22 wild type (n=3), TS22 muscleless (n =2), TS25 wild type (n=3), TS25 muscleless (n=1), TS27 wild type/muscleless (n=3).

### Mechanostimulation bioreactor culture of wild type and muscleless-limb rudiments

Our immunofluorescence data demonstrates that the establishment of a normal collagen network is dependent on skeletal muscles. The impact of the muscles could be due to 1) mechanical loading, 2) biochemical signalling from the muscles, or 3) a combination of the two. To test the hypothesis that the change in mechanical loading is the dominant factor affecting collagen organisation when skeletal muscle is absent, we performed bioreactor experiments in which limb explants underwent either *in vitro* dynamic mechanical stimulation, or static culture. Embryos were harvested at e15.5, which, in our hands, yields embryos at around TS24. From each embryo, one forelimb was cultured under dynamic loading conditions and the contralateral limb under static conditions, over a culture period of six days. Limbs were loaded three times per day, for two hours each time, under cyclical compression at 0.67 Hz. An applied displacement of 2 mm induced a flexion of approximately 14° at the elbow. Paired comparisons of the collagens most dramatically affected by the absence of muscles *in vivo* (collagens II, X and XI) were made between dynamically cultured and contralateral statically cultured forelimbs.

#### Dynamic loading in vitro affects localisation and structure of collagens II & X, but not XI

Dynamic loading promoted localisation of collagen II in the proximal future articular cartilage regions and in the distal epiphysis. In the dynamically cultured limbs, we consistently observed a distinct, uniform band of collagen II distribution in the future articular cartilage region of the humeral head (Figure 7A, C, dashed lines), as also seen *in vivo* at TS27 (Figure 2). In the proximal humerus of statically cultured limbs, collagen II localisation extended beyond the future articular cartilage region into the epiphysis (Figure 7B, D, dashed lines), leading to the loss of a distinct boundary between the future articular cartilage region and the rest of the humeral head. Dynamically cultured limbs had a stronger collagen II immunopositivity in the distal humerus compared to the statically cultured explants (Figure 7A–D, arrows). Dynamic culture also led to differences in collagen II structure in the proliferative region of the growth plate, but there was variation between the three samples analysed, as shown in Figure 7. In the dynamically cultured limbs of samples one and two (S1, S2), the collagen II organisation was similar to that normally seen at TS27 (Figure 2), with distinct longitudinal (Figure 7a, stars) and transverse septal organisation (Figure 7a, arrowheads). Localisation was more pronounced in the longitudinal septa than in the transverse septa, leading to a ‘rungs on a ladder-like’ organisation. In the statically cultured contralateral limbs of the same samples, a clear structure was present (Figure 7b), but there were no differences in collagen organisation between the longitudinal (Fig 7b, stars) and transverse septa (Fig 7b, arrowheads). In the third sample (S3), dynamic loading led to a structure which was not as pronounced as that of the other two dynamically cultured limbs (Fig 7c), and yet, a structured arrangement was still present with a prominent transverse septa (Fig 7c, arrowheads) and mild localisation in the longitudinal septa (Fig 7c, stars). In contrast, no septal organisation was observed in the cartilage of the statically cultured contralateral limb of S3, where collagen II was present throughout the matrix with large cell-shaped voids (Fig 7d). The variation in effects of loading on collagen II structure between S1/S2 and S3 is likely due to differences in the developmental stage of the animals, with sample three being more developmentally mature than the other two (as indicated by more pronounced morphology of the proximal and distal humerus in sample three (Supplementary Figure 9). These findings demonstrate the importance of mechanical loading for collagen II localisation and structure, and also suggest that the effects of mechanical stimulation on collagen II organisation depend on the rudiment developmental stage.

**Figure 7:**
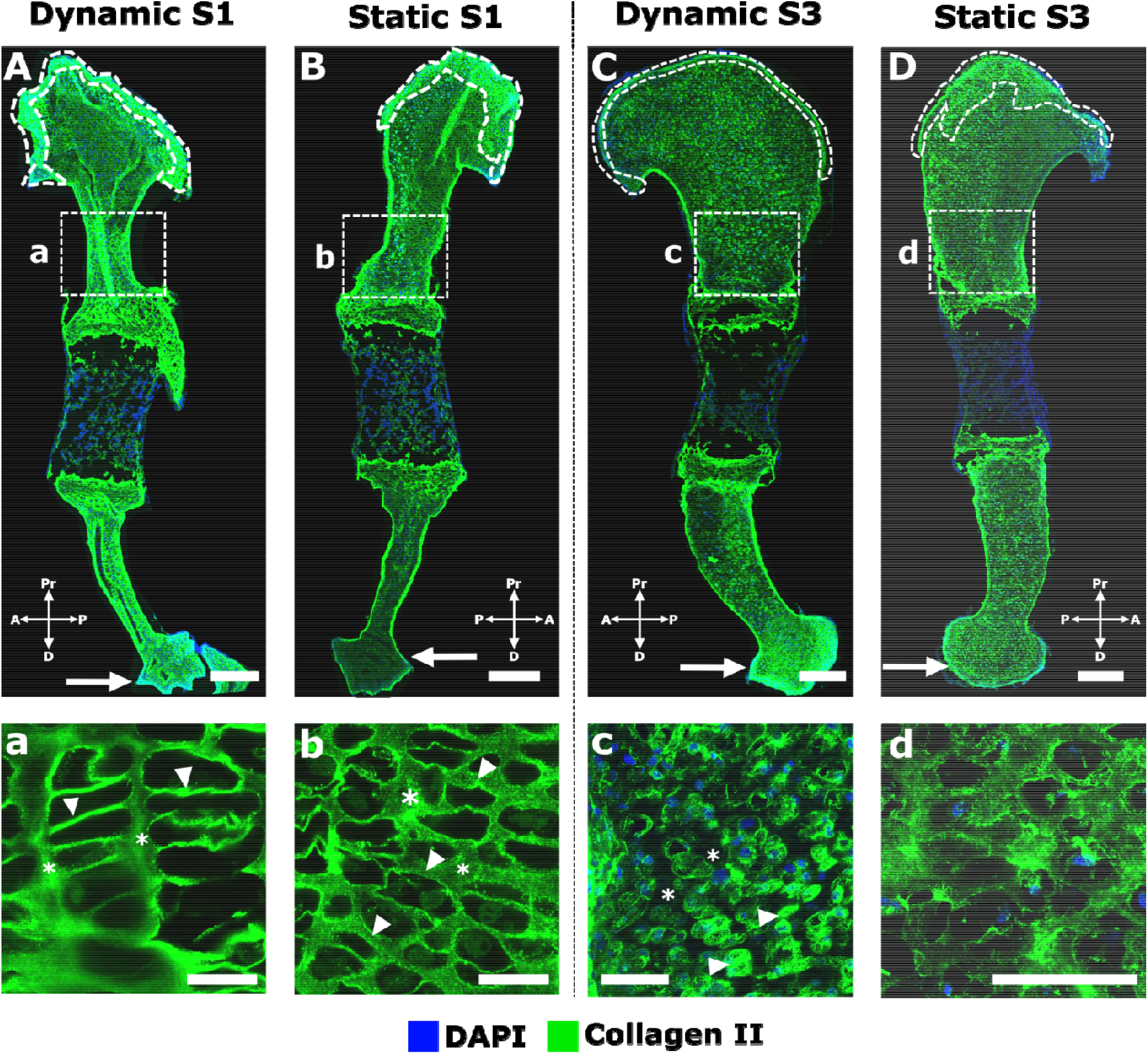
Collagen II localisation and structure were regulated by *in vitro* mechanical loading in wild type explants. A–D: Dynamic loading promoted localisation of collagen II in the proximal future articular cartilage regions and in the distal epiphysis; compare the distinct, uniform band of collagen II distribution in the future articular cartilage region of dynamically stimulated limbs (dashed lines in A, C) with the absence of a distinct boundary between the band of collagen II and the rest of the humeral head in contralateral statically cultured limbs (dashed lines in B, D). Collagen II immunopositivity was stronger in the distal humeri of dynamically stimulated limbs than in the statically cultured contralateral limbs (arrows in A–D). Dynamic loading *in vitro* also led to a more structured organisation of collagen II than seen in statically cultured contralateral limbs; compare the longitudinal septa (*) and transverse septa (arrowheads) between a & b, and c & d. Scale bars 200 µm (A–D), 20 µm (a–d). Sample numbers: n=3 embryos, right limbs cultured under dynamic loading and left limbs cultured under static conditions.

Dynamic loading *in vitro* led to more normal localisation of collagen X than static culture. As with collagen II, there was variation in the effects on collagen X between the limbs analysed, as shown in Figure 8. In the set of forelimbs from sample one (S1), dynamic loading led to localisation of collagen X to the growth plates (Figure 8A) and mineralising region (Fig 8A), while the statically cultured contralateral rudiment had extensive collagen X in the non-mineralised cartilage (Figure 8B). Therefore, while S1 did not have normal localisation in the dynamically cultured limb (due to immunopositivity in the mineralised cartilage), the localisation pattern was more normal than in the statically cultured limb. The strong immunopositivity observed in the non-mineralised cartilage of the statically cultured limb was characteristic of the muscleless-limb profile of collagen X at TS25 and TS27 (Figure 5). In the second pair of forelimbs (S2), after dynamic loading, collagen X was localised mainly to the growth plates (Figure 8C) with only mild immunopositivity in the mineralising cartilage (Figure 8C), while the static contralateral limb had strong immunopositivity in both the growth plate (Figure 9D) and the mineralising cartilage (Figure 8D). There were no obvious differences in collagen X structure in the growth plate in either sample (Figure 8a–d), with dynamic and static samples exhibiting a similar membrane-like configuration in this region similar to those seen in TS25 wild types (Figure 5). These results identify a role of mechanical loading for correct localisation of collagen X in the developing limb.

**Figure 8:**
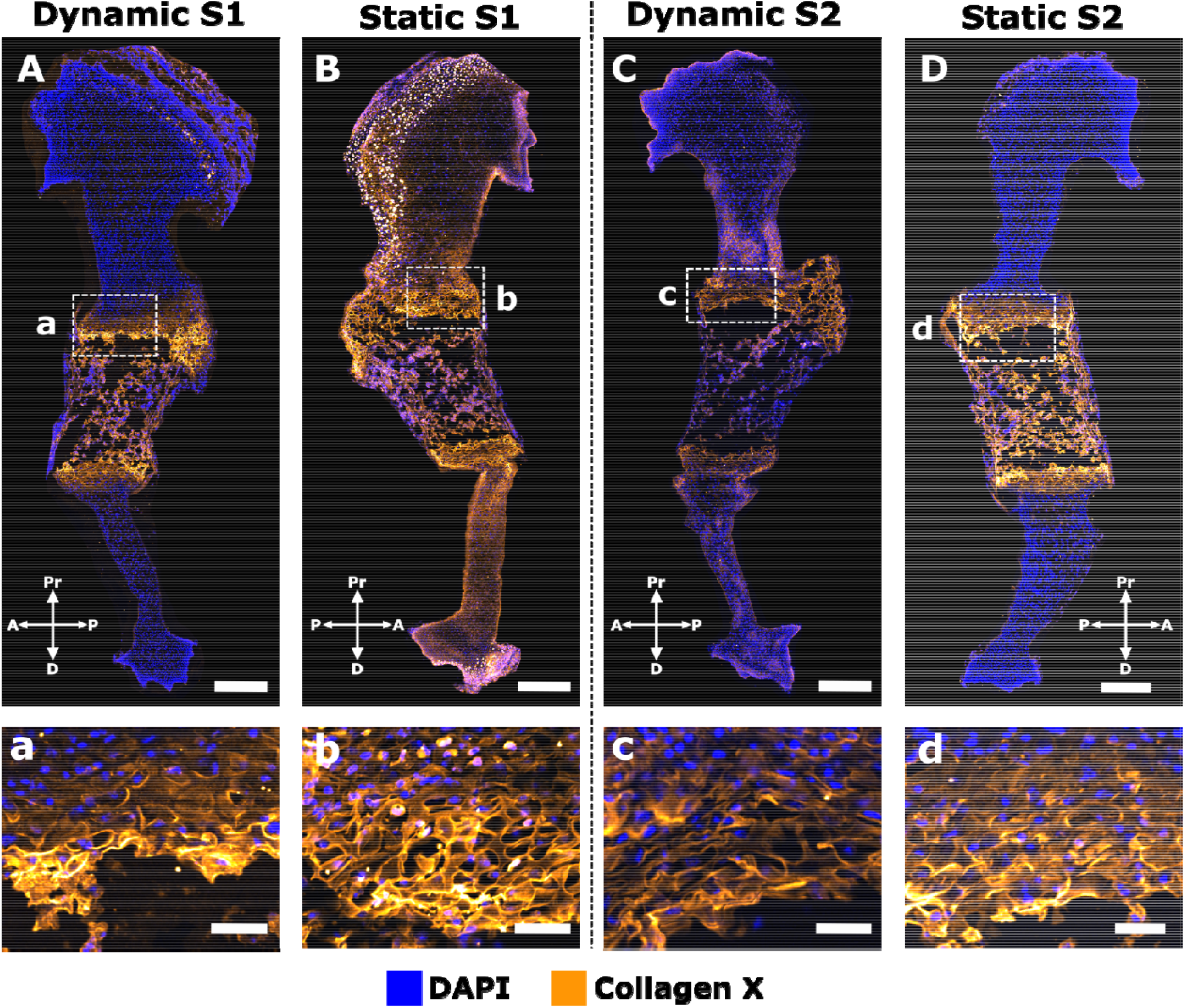
External mechanical loading determined localisation of collagen X in wild type explants. A–D: Dynamic loading *in vitro* led to more normal localisation of collagen X than static culture, with statically cultured limbs having more extensive collagen X in the non-mineralised cartilage (B) or in the mid-diaphysis (D). a–d: Dynamic loading *in vitro* had no obvious effects on the collagen X structure in the growth plate (. Scale bars 100 µm (A–D) and 20 µm (a–d). Sample numbers: n=2 embryos, right limbs cultured under dynamic loading, left limbs cultured under static conditions.

**Figure 9:**
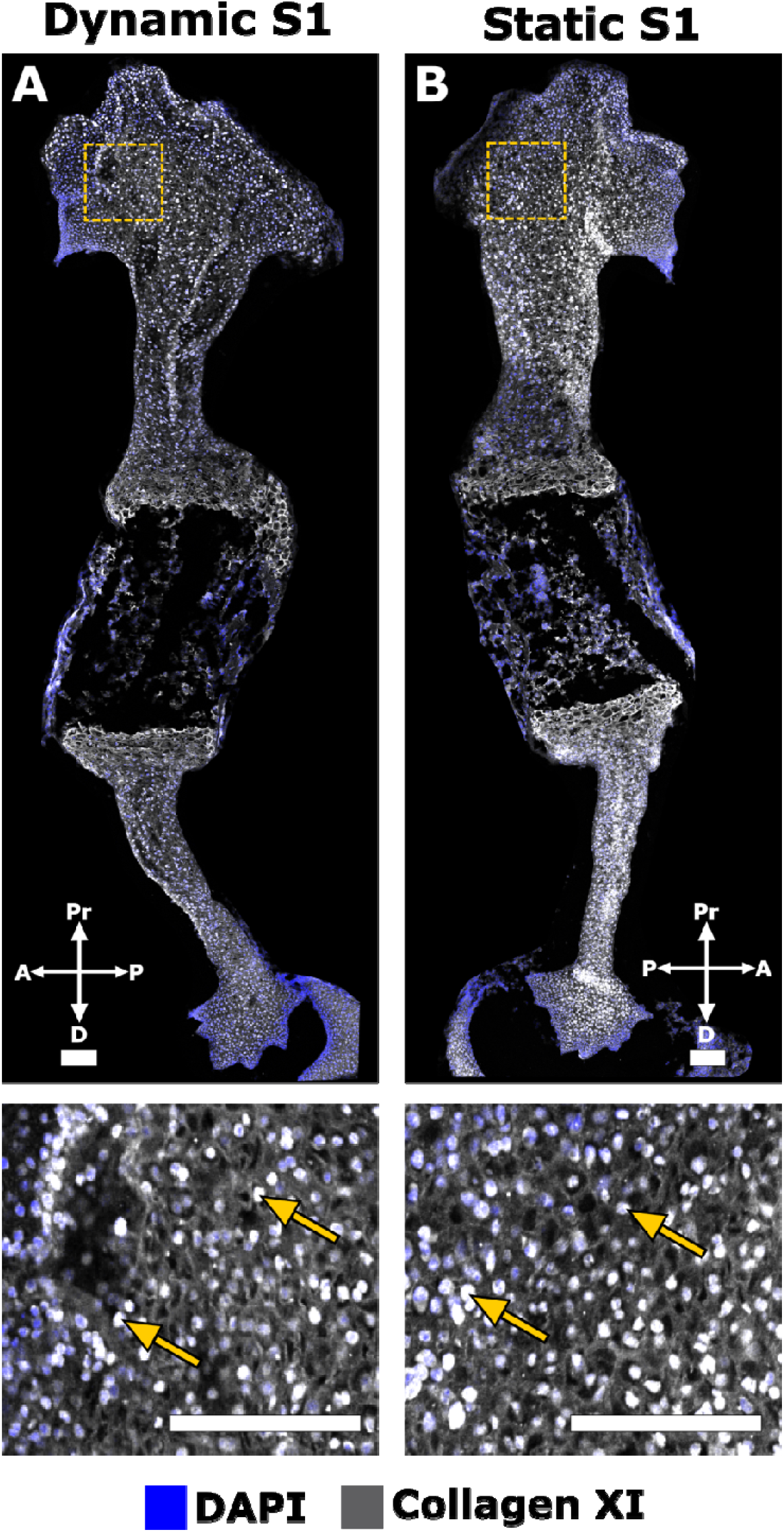
At the stage of development studied, external mechanical loading did not directly influence the structure or localisation of collagen XI in wild type explants. No difference in collagen XI organisation was observed between dynamically (A) and statically (B) cultured limbs harvested at e15.5. There was widespread intracellular localisation of collagen XI in both dynamically and statically cultured limbs (arrows). Scale bars 100 µm. Sample numbers: n=2 embryos, right limbs cultured under dynamic loading, left limbs cultured under static conditions.

Neither the structure nor the distribution of collagen XI were different between dynamic and static limbs in either of the two samples examined (Figure 9). An indistinct mesh-like organisation was observed throughout the rudiment in both dynamic and static limbs (Figure 9). This pattern of localisation and structure was not similar to any stages of the normal wild type samples (Figure 6). Prominent intracellular localisation was observed in both sets of limbs (arrows in Figure 9), as seen in the muscleless limbs at TS27 (Figure 6). As collagen XI was not substantially affected by absent skeletal muscle *in vivo* until e18.5 (TS27, Figure 6), it is possible that the rate of development and/or duration of culture were insufficient for effects of loading on the collagen XI network to become evident.

#### Dynamic mechanical loading in vitro rescues aspects of collagen network emergence in muscleless-limb explants

Our final experiment sought to investigate if *in vitro* mechanical loading could reverse or recover the adverse effects of fetal immobility on development of collagens II, X and XI, further corroborating a direct link between mechanical loading and the collagen network development, and identifying a possible therapeutic intervention for healthier skeletal development after a period of fetal akinesia. Muscleless-limbs were harvested at e15.5, or approximately TS24, at which stage collagens II and X were already abnormal in the mutants (Figure 2 and 5), then cultured under static or dynamic conditions.

Aberrant retention of collagen II in the mineralised region was seen in both statically and dynamically cultured muscleless-limbs (n=3, Figure 10A, B), indicating that abnormal localisation of collagen II in muscleless-limbs at time of harvest could not be recovered by mechanical stimulation *in vitro*. Strong localisation of collagen II to the future articular cartilage region was observed in the dynamically cultured, but not the statically cultured, muscleless-limbs (compare Figure 10B and Figure 10A, dotted lines), identifying a direct role of mechanical loading in specification of collagen II localisation in the future articular cartilage region of the proximal humerus. With regards to the structure of collagen II (also aberrant in the muscleless-limbs at time of harvest), dynamic loading rescued aspects of structure in the proliferative region of the growth plate. In the dynamically cultured limbs, distinct longitudinal (Figure 10b, arrowheads) and transverse septal (Figure 10b) arrangements were evident, with the transverse septa surrounding cell-shaped voids in a columnar arrangement.. The structure in the dynamically cultured limbs showed some resemblance to the structure seen in the TS25 wild type rudiments (Figure 2e). In the statically cultured contralateral limbs, no distinct longitudinal or transverse septal organisations of collagen II were observed (Figure 10a). The thick bundles of collagen II in the static group lacked any obvious principle arrangement (Figure 10a, arrowheads) and surrounded randomly-oriented cell-shaped voids. Taken together, these data indicate that mechanical loading is critical to restricted localisation of collagen II in the future articular cartilage region and to organisation of collagen II in the proliferative region of the growth plate.

**Figure 10:**
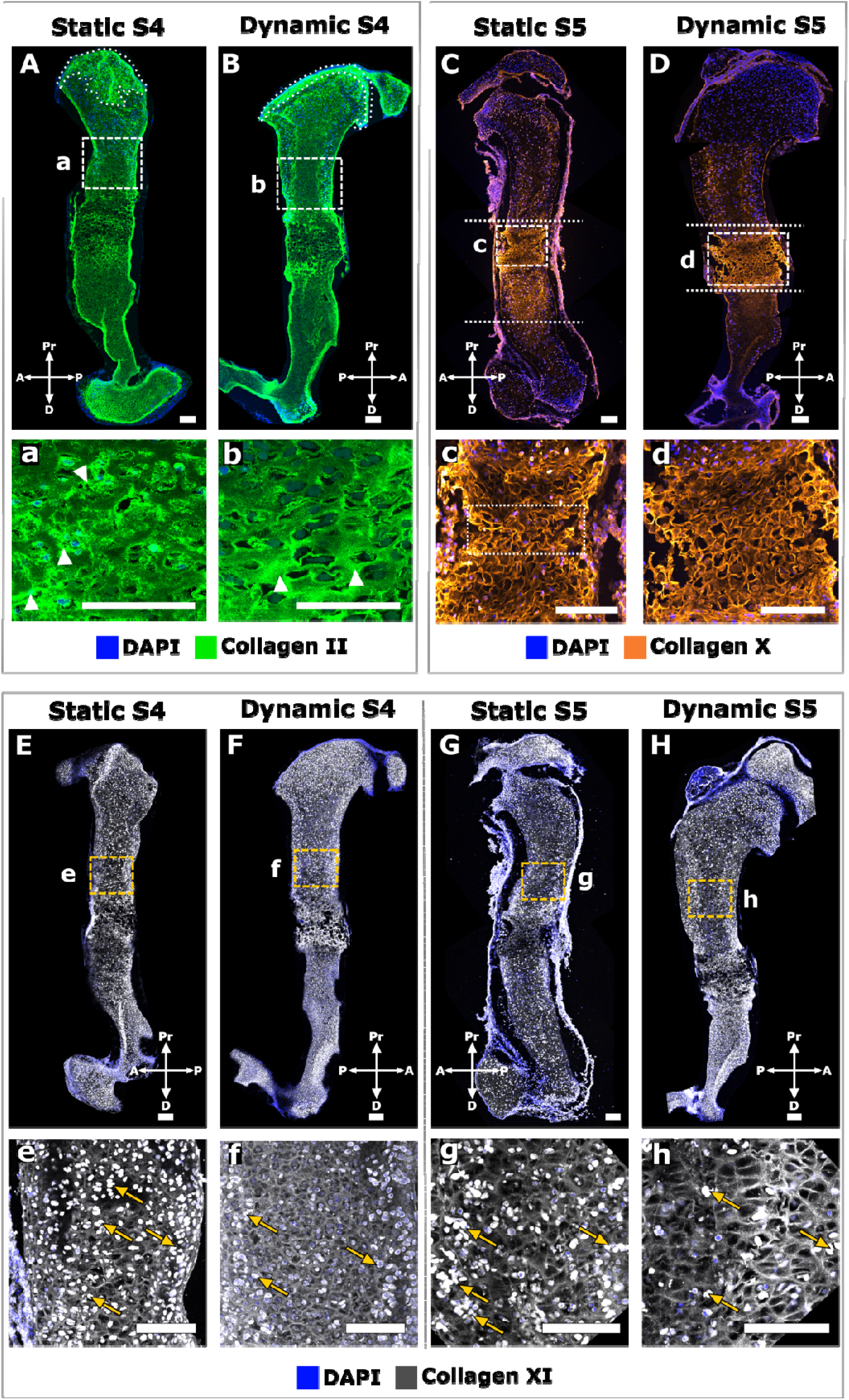
External mechanical loading partially restored the localisation and structure of collagens II and X and had subtle effects on the localisation and structure of collagen XI in cultured muscleless-limb explants. A, B, a, b : *In vitro* loading of muscleless-limb explants restored more normal localisation of collagen II in the future articular cartilage region (dotted areas in A, B) and rescued aspects of structure in the proliferative region of the growth plate, with distinct longitudinal and transverse septal arrangements evident in the dynamically cultured limbs (compare the longitudinal septa (arrowheads) and transverse septa between a, b). C, D, c, d: *In vitro* loading of muscleless-limb explants promoted mid-diaphyseal localisation of collagen X, with reduced and milder immunopositivity in the diaphysis and epiphyses (D) compared to the statically cultured contralateral limbs (C). Dynamic loading also promoted more normal collagen X structure in the muscleless-limb explants, with an intricate interlocking collagen X matrix organisation without any collapsing regions in the loaded limbs (c, d,). E–H, e–h: Loading of muscleless-limbs *in vitro* led to a more pronounced collagen XI meshwork pattern (compare within contralateral limb pairs shown in E–H), and a reduction in intracellular localisation compared to the statically cultured muscleless-limbs (arrows in e–h). Scale bars 100 µm. Sample numbers: n=3 embryos for collagens II and X, n=2 for collagen XI, left limbs cultured under dynamic loading and right limbs cultured under static conditions.

*In vitro* mechanical loading partially rescued the effects of fetal immobility on the localisation and structure of collagen X in muscleless-limb explants. Punctate collagen X localisation was present outside of the mineralising regions in both static and dynamic muscleless-limb explants (n=3, Figure 10C, D), as seen in muscleless-limbs *in vivo* and in statically cultured wild type explants. However, dynamic loading *in vitro* promoted mid-diaphyseal localisation of collagen X, with reduced and milder immunopositivity in the diaphysis and epiphyses (Figure 10D) compared to the statically cultured contralateral limbs (Figure 10C). Dynamic loading also promoted a more normal collagen X structure in the muscleless-limb explants. In the static group, collagen X had a membrane-like configuration with regions where the matrix had a collapsed appearance (Figure 10c, dotted box). With dynamic loading, an intricate interlocking collagen X matrix organisation without any collapsing regions was restored (Figure 10d), somewhat resembling the oblique/teardrop appearance seen in the TS25 rudiments (Figure 5e). Therefore, these data corroborate a direct link between mechanical loading and both localisation and structure of collagen X.

As the effects of fetal immobility on collagen XI did not become evident until TS27 (Figure 6), substantially after the time of harvest for culture (e15.5/TS24), the concept of rescue or recovery due to loading *in vitro* did therefore not apply for collagen XI. However, assessing the effects of external mechanical stimulation on collagen XI in the muscleless-limbs enabled us to test if the early establishment of the collagen XI network occurs completely independent of mechanical loading. Loading of muscleless-limbs *in vitro* led to a more pronounced collagen XI meshwork pattern (compare within contralateral limb pairs shown in Figure 10E–H), and a reduction in intracellular localisation compared to the statically cultured muscleless-limbs (arrows, Figure 10e– h). These results are in contrast to the same experiment performed on limb explants with normal muscle, which showed no effects of *in vitro* loading on collagen XI (Figure 9). Therefore, the influence of *in vitro* mechanical loading on the collagen XI network depends on the *in vivo* loading history of the cultured limbs. We conclude therefore that mechanical loading from skeletal muscle contractions is likely to contribute to early events in establishment of the collagen XI network, even though the effects of fetal immobility on collagen XI do not become apparent until TS27.

## Discussion

This study provides evidence for the role of muscle contraction-induced mechanical stimulation in promoting the correct spatial localisation and structure of key cartilage and bone collagens over prenatal skeletal development. In the Splotch delayed (“muscleless-limb”) model, spatial localisations of collagens I, II, V and X were aberrant, as were the structural organisations of collagens I, II, VI, X and XI. Effects on localisation and structure varied over the three stages studied, with differences in localisation being particularly pronounced at the earliest stage (TS22) and differences in structure becoming more pronounced with advancing development. We next tested the hypothesis that the effects of absent skeletal muscle on collagen structure and localisation are predominantly due to the absence of muscle contraction induced mechanical loading. Mechanical loading *in vitro* directly modulated the structure and localisation of collagen II and the localisation of collagen X. Collagen XI was not directly modulated by *in vitro* loading, which may be related to the age at which the limb explants were harvested. Finally, we investigated if mechanical loading *in vitro* could rescue any of the effects of fetal akinesia on collagen network emergence by culturing muscleless-limb explants. Mechanical loading *in vitro* rescued some features of the collagen network seen in the muscleless-limbs, including collagen II localisation in the future articular cartilage region of the proximal humerus, collagen II structure in the proliferative region of the growth plate, and localisation and structure of collagen X. We conclude therefore that skeletal muscle is essential for the normal development of the key cartilage and bone collagens and that mechanical loading from muscle contractions is directly involved in the establishment of the collagen II and X networks.

This work has shown that skeletal muscles are critical for the correct structural organisation of key collagens in prenatal cartilage and bone. The importance of mechanical loading for collagen fibre organisation has been previously demonstrated in postnatal and adult articular cartilage (*5, 14, 30-35*). Collagen fibres align perpendicular to the loading direction in tissue-engineered cartilage constructs grown under unconfined compression (*32-34*). Brama et al., 2009 (*5*) showed that high-intensity exercise in horses led to increased collagen parallelism and affected collagen orientation, with region-specific effects according to loading of individual joint sites. In hamsters, wheel-exercise led to altered collagen network organisation and collagen content compared to non-exercised controls (*14*). Changes in collagen content or organisation, were also found in wheel-exercised guinea pigs (*31*), while in dogs, alterations in collagen orientation but no changes in collagen content were found after an intense exercise programme (*30*). The advance of the current study is that it demonstrates that skeletal muscles are essential for collagen fibre localisation and structure in prenatal cartilage and bone, and also identifies which of the major skeletal collagen types are most- and least-sensitive to muscle loading.

Time-variant effects of muscle or loading were apparent for some collagens, particularly collagens II and XI. *In vivo*, absent skeletal muscle was associated with precocious formation of the collagen II mesh-like structure at early stages, which was lost by the latest stage examined. Our hypothesis is that muscle loading is a constraining factor for the early formation of collagen II structure, but that loading is needed to sustain healthy development of the network. Collagen XI was dramatically affected by absent muscle *in vivo* only at the latest stage of development studied, while collagen XI in limbs harvested at e15.5 was unaffected by mechanical loading *in vitro*. These data would indicate either that early establishment of the collagen XI network is independent of mechanical loading, or that the effects of absent loading do not become apparent until a later developmental stage. Our finding that the collagen XI network was more normal in mechanically stimulated muscleless-limbs compared to statically cultured muscleless-limbs indicates that the latter hypothesis is more likely, that early loading (prior to e15.5) of the limbs contribute to later collagen XI emergence.

Different levels of sensitivity to loading were exhibited in different regions of the developing rudiment, particularly *in vitro*. The effects of *in vitro* mechanical loading on collagens II and X were pronounced in the growth plate and future articular cartilage regions, but not in the epiphyseal (resting) cartilage. *In vitro* cultured limbs were harvested at e15.5, by which time the key events of epiphyseal cartilage collagen development have already initiated. We propose therefore that early stimulation of the limbs (prior to e15.5) is most important for initiation of the appropriate collagen network in the epiphyseal cartilage. In contrast to the epiphyseal cartilage, the prenatal growth plate requires sustained mechanical stimulation for proper development of its collagens, likely due to the dynamism of its development (*36, 37*). The future articular cartilage region may also be undergoing dynamic processes in the chosen timeframe based on the observed changes in collagen II localisation in the region between statically cultured and mechanically stimulated limbs. Mechanical loading has not yet been implicated in prenatal development of the future articular cartilage, despite the known importance of loading for postnatal articular cartilage (*5, 14, 30-35*), providing substantial scope for further investigation of the role of mechanical loading in directing articular cartilage development.

Our study is not without limitations. Tissue sections were used to characterise the collagen distribution and network architecture and, therefore, the 3D organisations were not characterised. Combining tissue clearing techniques for immunofluorescence with light sheet microscopy would enable characterisation of collagens in greater resolution and through a greater depth of the rudiment. Given that mechanoregulated activity has been associated with collagens III (*38*), a fibril network modifier which stabilises the collagen II network (*39, 40*), a lack of alteration in collagen III when skeletal muscle was absent is surprising. It is possible that immunofluorescence is unsuitable for detecting the architecture of collagen III in sufficient details. Future studies could analyse collagen interactions by attenuated total reflection-Fourier transform infrared (ATR-FTIR) spectroscopy in order to verify if collagen III develops independently of skeletal muscles. Finally, sample numbers for all groups were such that statistical analyses were not feasible. However, studying multiple developmental stages, and both *ex vivo* and *in vitro* loading scenarios, strengthens our conclusions.

The work presented here has provided novel insights into the mechanoregulation of the increasing complexity of the multiscale collagen network over skeletal development. We have demonstrated a direct role for mechanical loading for emergence of collagen distribution and structure during prenatal skeletal development. We have shown that collagens I, II, V, VI, X and XI are sensitive to the absence of skeletal muscles *in vivo*, and that collagens II and X are directly modulated by mechanical loading *in vitro*. These findings further our understanding of cartilage and bone matrix development, and in particular the effects of fetal akinesia on the collagen matrix of skeletal tissues. The finding that external mechanical loading can partially rescue the effects of fetal akinesia on the collagen matrix opens avenues towards physical therapies which could mitigate against the effects of reduced or abnormal fetal movements on skeletal development. Furthermore, the key to successful regeneration of bone and cartilage (which has so far remained elusive (*41*) may be through understanding how the complexity arises over development, i.e., a developmentally-inspired approach. The advance in understanding on the contribution of mechanical loading to collagen network establishment and maturation provided by this research will facilitate future advances in developmentally-inspired replication of the collagen architecture in engineered skeletal tissues.

## Materials and Methods

### Tissue collection and processing

All experiments were performed in accordance with European legislation (Directive 2010/63/EU). For the immunofluorescence of collagens, and for the cultures of muscleless limbs, heterozygous *Splotch-delayed* (Pax3^spd/+^) females and males (imported from The Jackson Laboratory, Maine, USA; JAX stock #000565) were crossed to generate homozygous Pax3^Spd/Spd^ ‘muscleless limb’ embryos. For limbs designated for immunofluorescence of collagens, embryos were harvested at embryonic days 13.5, 16.5 and 18.5 and selected for analysis based on being categorised into Theiler stages TS22, TS25 or TS27 (*16*). The wild type limbs used for immunofluorescence of collagens were the same limbs described in our previous study (*9*). For the wild type explant cultures, embryos were harvested from pregnant C56Bl/6J mice (Charles River (Massachusetts, USA; stock # 3062894) at embryonic day (e) 15.5. For the muscleless limb cultures, Pax3^Spd/Spd^ muscleless limb embryos were harvested at e15.5. All embryos from the *Splotch-delayed* line were genotyped using PCR on DNA derived from head tissue. The PCR reaction was carried out for 30 cycles, each with duration of 30 seconds at 94 °C, 60 °C and 74°C, using three primers. The primer sequences used were AGGGCCGAGTCAACCAGCACG & CACGCGAAGCTGGCGAGAAATG for controls and AGTGTCCACCCCTCTTGGCCTCGGCCGAGTCAACCAGGTCC & CACGCGAAGCTGGCGAGAAATG for mutants. Sample numbers are detailed in Table 1 and 2.

**Table 1:**
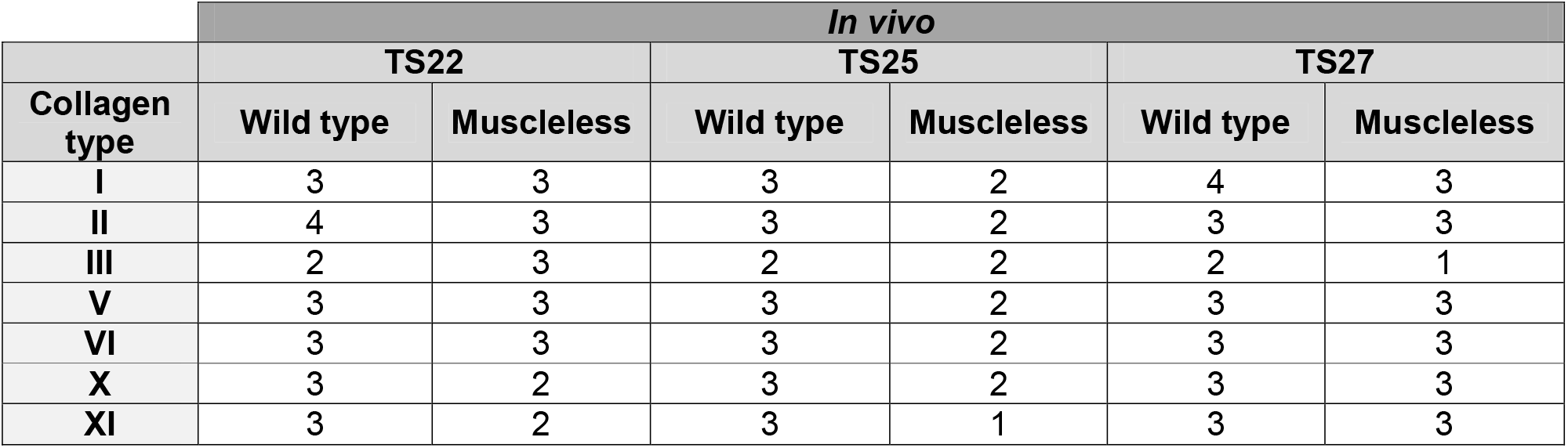
Numbers of limbs analysed for *in vivo* collagen characterisation experiments.

| Collagen type | *In vivo* |  |  |  |  |  |
| --- | --- | --- | --- | --- | --- | --- |
|  | TS22 |  | TS25 |  | TS27 |  |
|  | Wild type | Muscleless | Wild type | Muscleless | Wild type | Muscleless |
| I | 3 | 3 | 3 | 2 | 4 | 3 |
| II | 4 | 3 | 3 | 2 | 3 | 3 |
| III | 2 | 3 | 2 | 2 | 2 | 1 |
| V | 3 | 3 | 3 | 2 | 3 | 3 |
| VI | 3 | 3 | 3 | 2 | 3 | 3 |
| X | 3 | 2 | 3 | 2 | 3 | 3 |
| XI | 3 | 2 | 3 | 1 | 3 | 3 |

**Table 2:** Numbers of limbs analysed for *in vitro* studies

| Collagen type | Bioreactor |  |  |  |
| --- | --- | --- | --- | --- |
|  | Wild type |  | Muscleless |  |
|  | Static | Dynamic | Static | Dynamic |
| II | 3 | 3 | 3 | 3 |
| X | 2 | 2 | 3 | 3 |
| XI | 2 | 2 | 2 | 2 |

**Table 3:** List of antibodies used and their product details

| Antibody | Supplier | Catalogue No. |
| --- | --- | --- |
| Collagen I | Abcam, UK | ab34710 |
| Collagen II | Sigma Aldrich, UK | MAB8887 |
| Collagen III | Abcam, UK | ab7778 |
| Collagen V | Abcam, UK | ab7046 |
| Collagen VI | Abcam, UK | ab6588 |
| Collagen X | Abcam, UK | ab58632 |
| Collagen XI | Invitrogen, UK | PA5-77258 |
| Goat anti-rabbit (Cy3®) | Abcam, UK | ab6939 |
| Rabbit anti-mouse (Alexa Fluor® 488) | Abcam, UK | ab150125 |

### Immunofluorescence and image acquisition by confocal microscopy

Forelimbs were harvested and processed for cryosectioning. For cryoprotection, limbs were exposed to an increasing sucrose gradient (15% and 30% sucrose) and then embedded in OCT (optimal cutting temperature) (Agar Scientific, Stansted, UK) diluted with 50% sucrose. Twelve μm thick sections were cut using a cryostat (NX70, Leica Biosystems, UK) (*9*). For immunofluorescence, cryosections were permeabilised with 0.1% Tween 20 (Sigma-Aldrich)/1% dimethyl sulfoxide (DMSO; Sigma-Aldrich) in phosphate-buffered saline (PBS) and blocked with 5% (v/v) normal goat serum (Sigma-Aldrich). Following blocking, tissue sections were either treated with 10 mg/ml bovine testicular hyaluronidase (HA) (Sigma-Aldrich) for one hour (collagens III and XI) prior to incubation with primary antibody or directly incubated with primary antibodies without HA treatment (collagens I, II, V, VI and X). Primary antibody against specific collagen types were used in 1:50 dilutions. Following an eighteen hour incubation with the primary antibody at 4°C, tissues were washed and further incubated with a secondary antibody (1:200 dilutions) and 4′,6-diamidino-2-phenylindole (DAPI, SigmaAldrich) (1:2000 dilution) for two hours at room temperature. Details of both the primary and secondary antibodies are provided in Table 2. Direct fluorescence acquisition of labelled tissue sections was performed using an inverted confocal laser scanning microscope (Zeiss LSM 510 and Leica CF6) as described previously (*9*).

### Image analysis

Details of the number of limbs analysed for each experiment type are given in Tables 1 and 2. Confocal images were processed in FIJI (*17*). Whole rudiment images were created using Stitching (Pairwise stitching) plugin (*42*) in FIJI. Linear Blending was used as the method of Fusion and all channels were averaged for image registration. Where automatic stitching created blurry images in the overlapped zone, images were manually stitched in Inkscape (Inkscape Project, 2021, retrieved from https://inkscape.org).

### Mechanostimulation bioreactor protocol

Following harvest at e15.5, the forelimbs were finely dissected and the soft tissues were manually removed with the forceps (*43*). The forelimbs were then pinned at the scapula to a deformable foam base cut into rectangular steps as described previously (*44*). Limbs designated for dynamic culture were then placed within the Ebers TC-3 bioreactor chambers (Don Whitley Scientific, Bingley, UK) to undergo uniaxial compression. Contralateral limbs designated for static culture were placed (with the same foam support as the dynamic cultures) in a petri dish. Limb explants were cultured at an air-liquid interface using basal media (α-MEM GlutaMAX, Thermo Fisher Scientific, USA) supplemented with 1% penicillin-streptomycin with amophotericin B, 100µM ascorbic acid (Sigma Aldrich, USA), 2mM β-glycerophosphate and 100nM dexamethasone. Explant cultures were incubated for six days at 37°C in a humidified atmosphere containing 5% CO_2_ and the media was changed every 24 hours. Limb explants designated for dynamic culture were subjected to mechanical deformation. The loading regime comprised cyclical compression of the forelimbs at 0.67 Hz to a displacement of 2 mm for two hours, three times per day. This produced approximately 14° angle cyclic flexion of the elbow. The angle was estimated based on previous studies carried out on chick joints using the same experimental set-up (*44*). Paired comparisons of collagens were performed between the mechanically loaded forelimb and the contralateral statically cultured forelimb, from the same embryo.

## Notes

### Competing Interest Statement

The authors have declared no competing interest.

