## Supplementary Figures File for "Mechanical Loading due to Muscle Movement Regulates Establishment of the Collagen Network in the Developing Murine Skeleton"

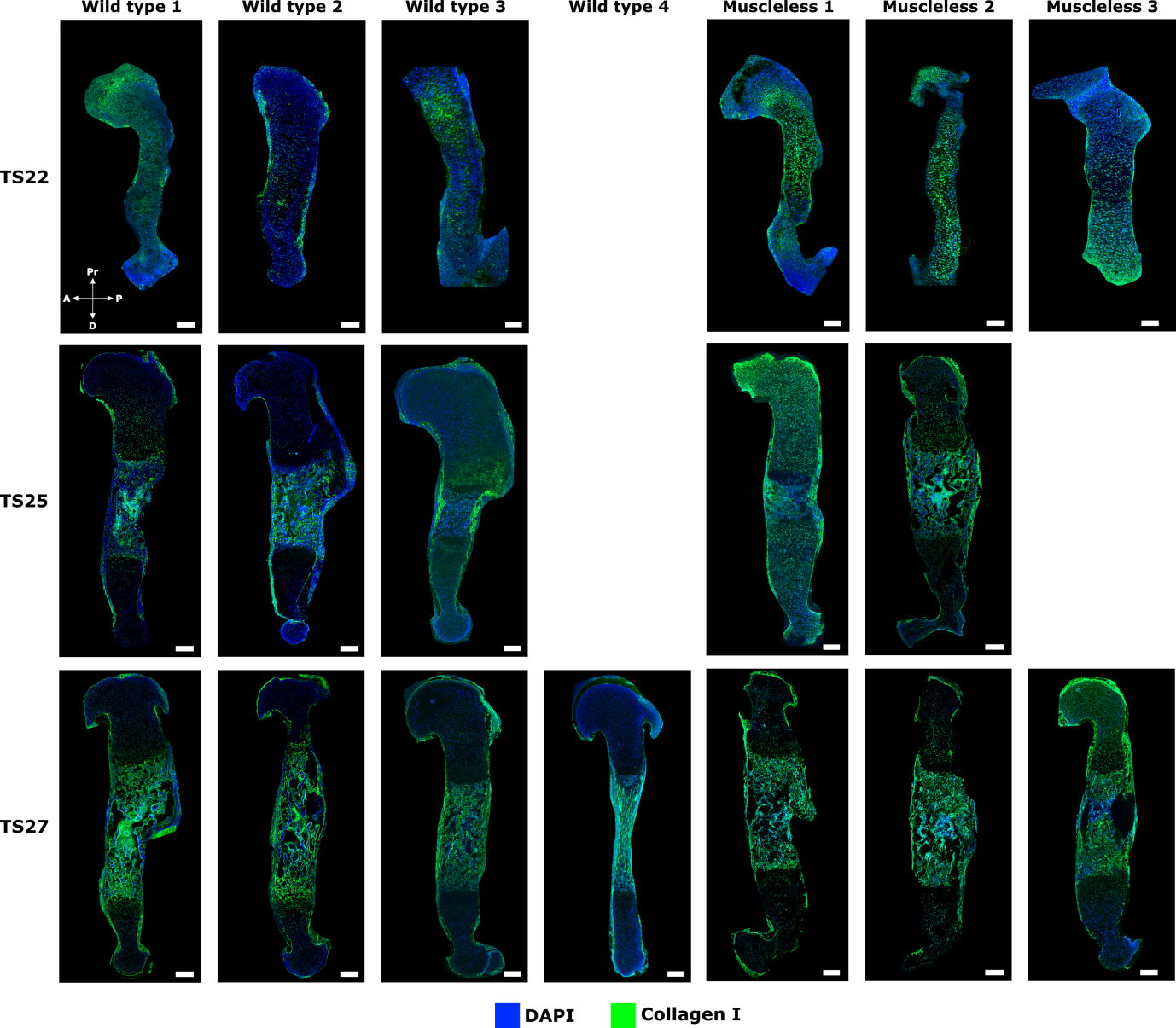

**Figure 1:** Immunofluorescence images of TS22, TS25 and TS27 wild type and muscleless limb humerus stained with collagen I antibody. Scale bars represent 200 μm.

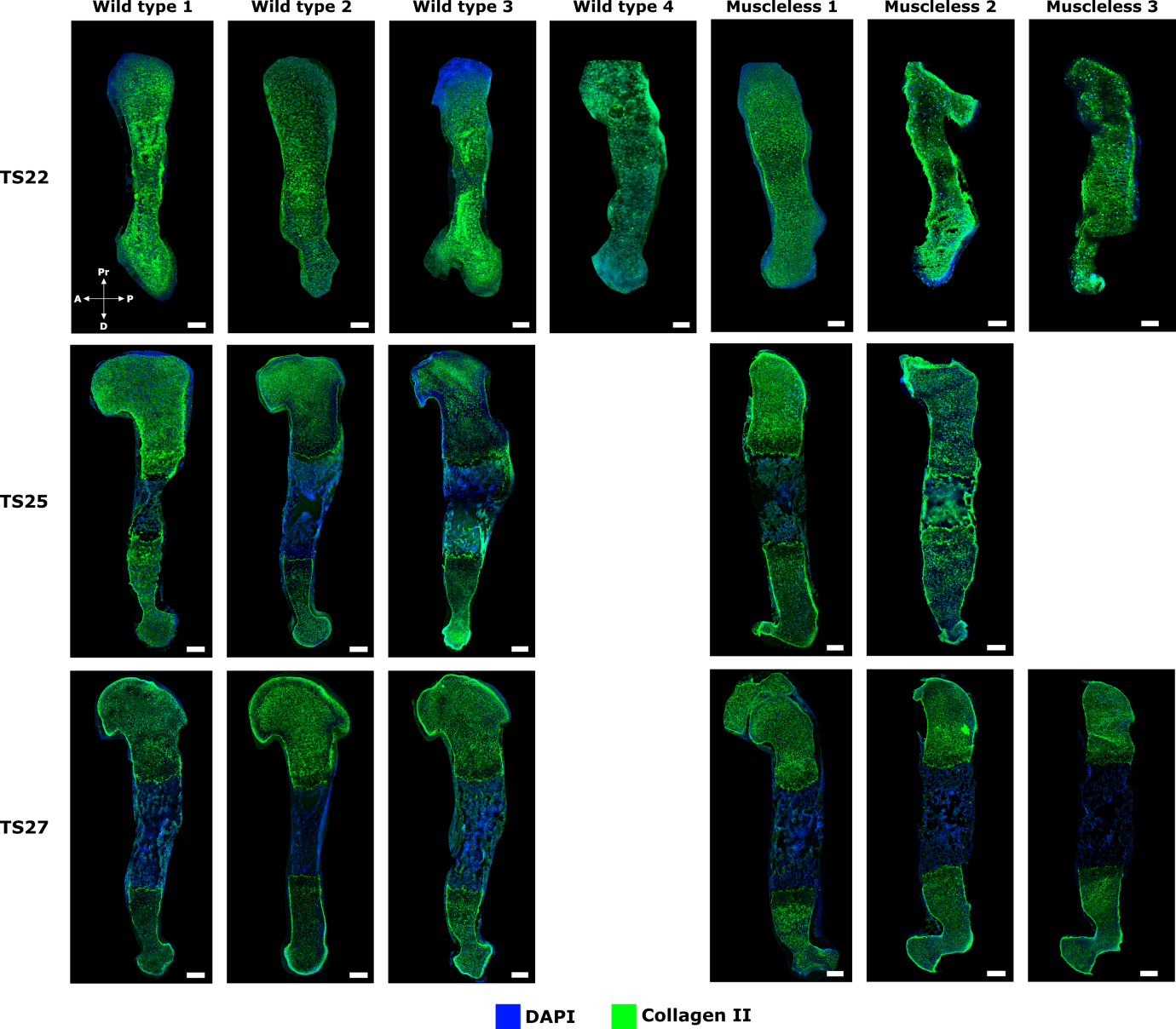

**Figure 2:** Immunofluorescence images of TS22, TS25 and TS27 wild type and muscleless limb humerus stained with collagen II antibody. Scale bars represent 200 μm.

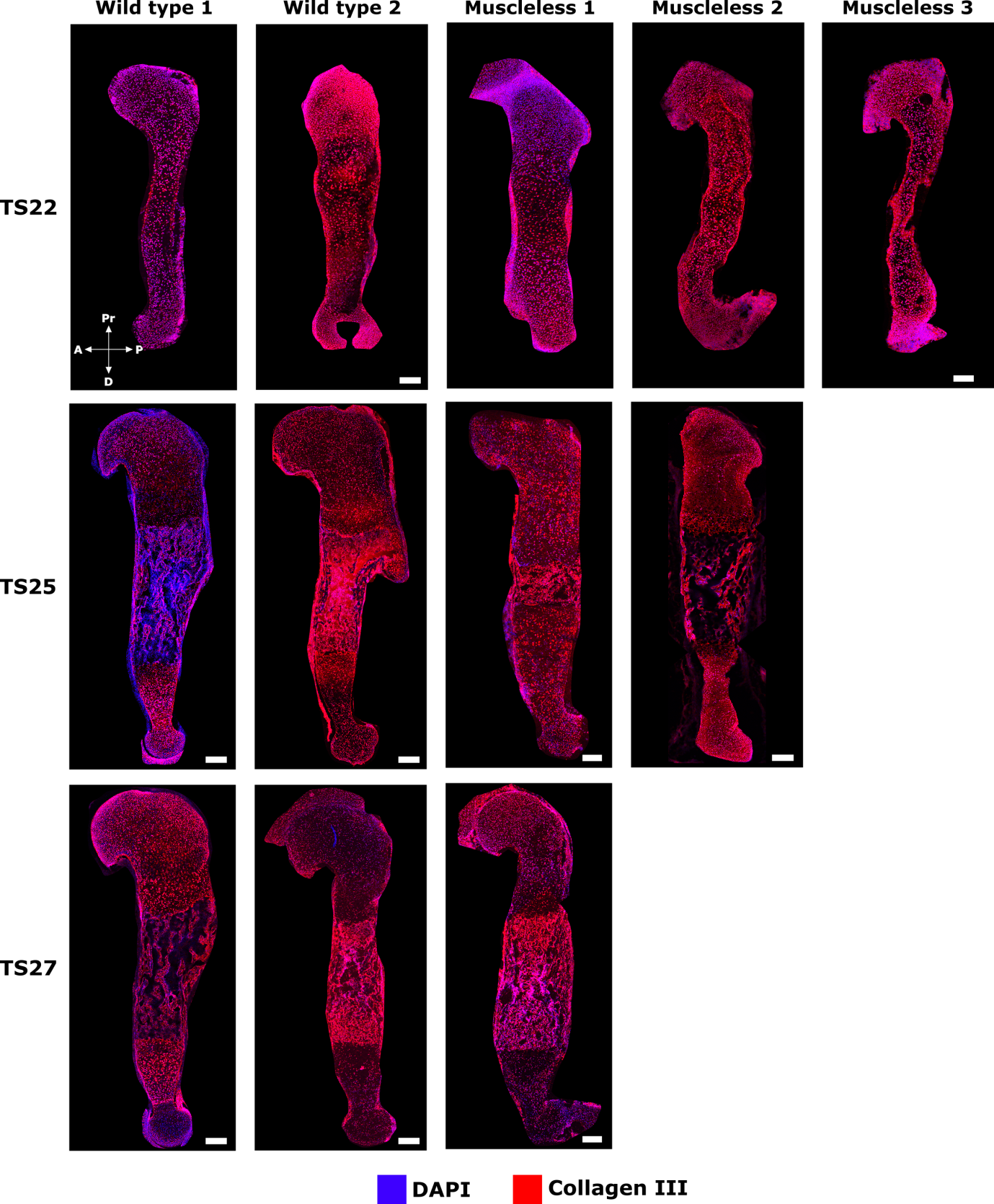

**Figure 3:** Immunofluorescence images of TS22, TS25 and TS27 wild type and muscleless limb humerus stained with collagen III antibody. Scale bars represent 200 μm.

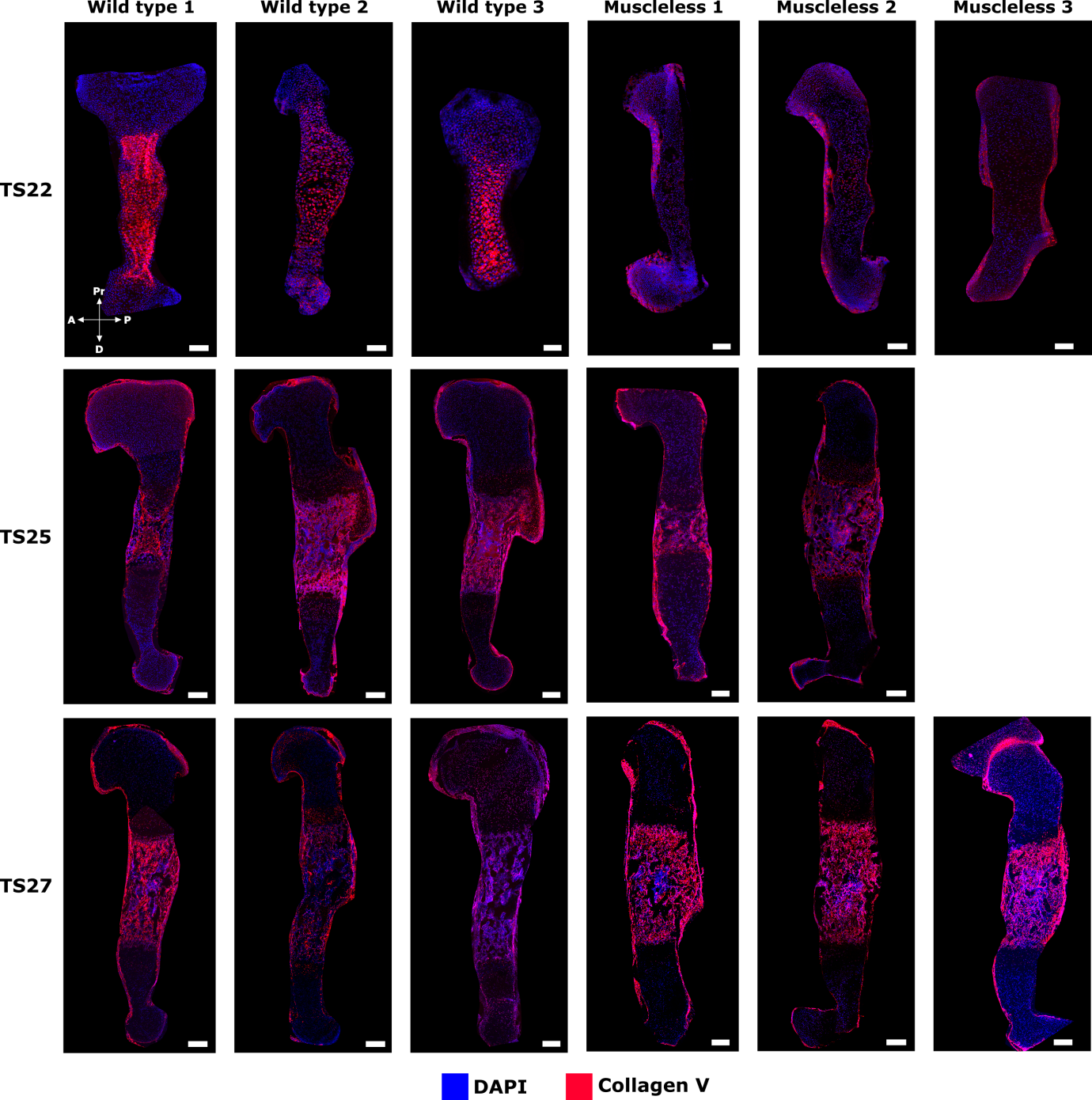

**Figure 4:** Immunofluorescence images of TS22, TS25 and TS27 wild type and muscleless limb humerus stained with collagen V antibody. Scale bars represent 200 μm.

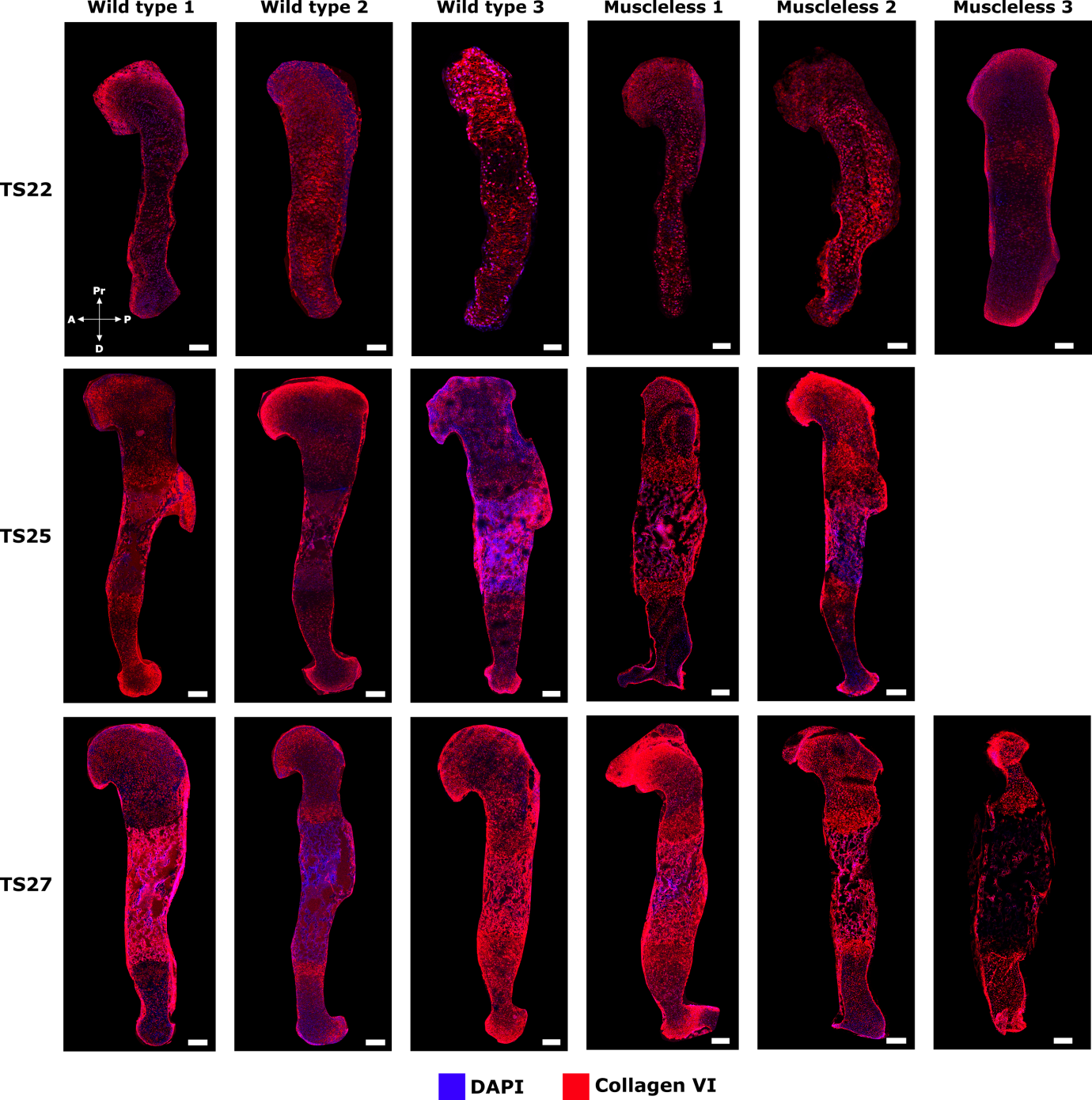

**Figure 5:** Immunofluorescence images of TS22, TS25 and TS27 wild type and muscleless limb humerus stained with collagen VI antibody. Scale bars represent 200 μm.

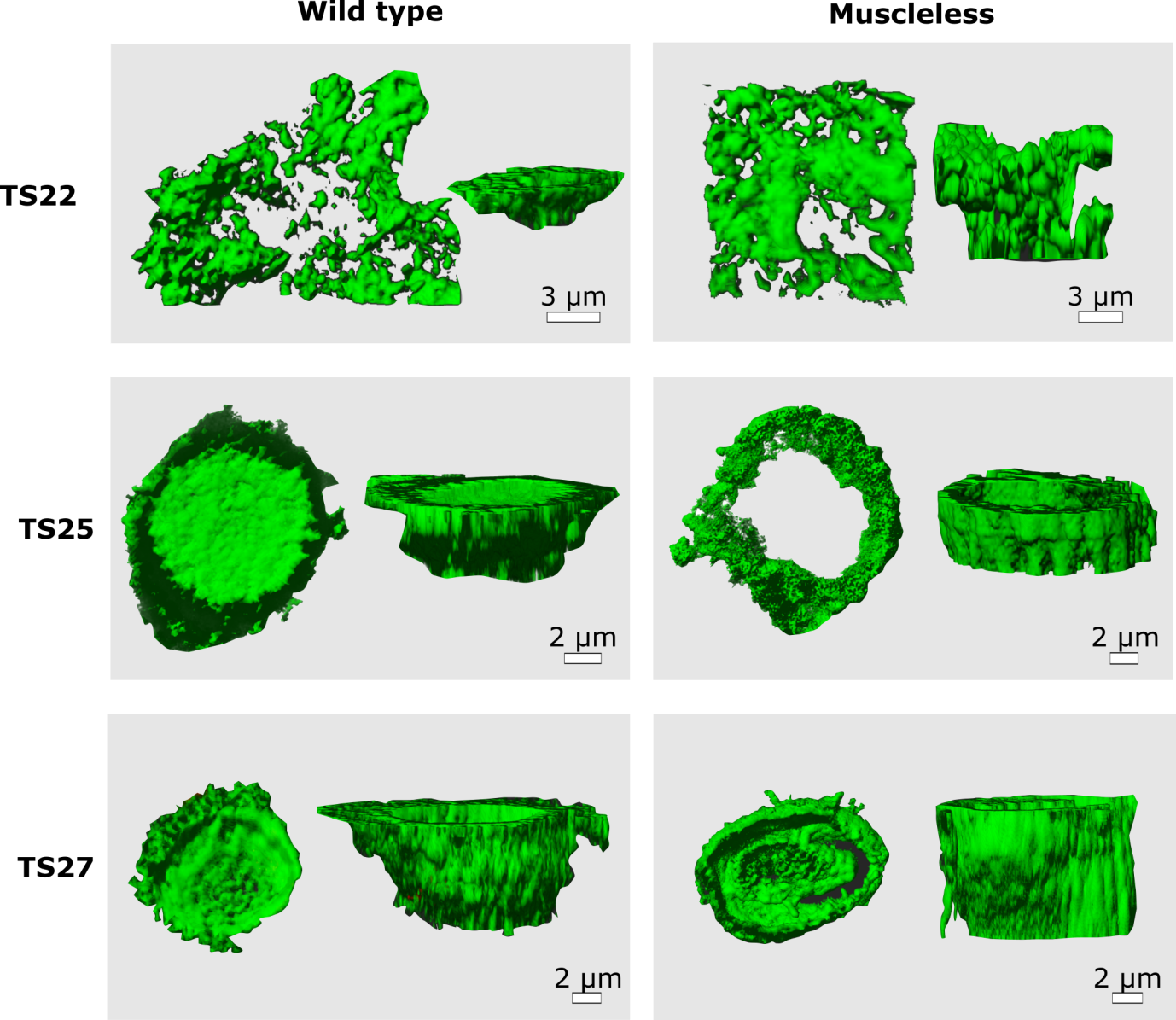

**Figure 6:** 3D representations demonstrating collagen VI containing chondron morphology at the humeral head region of wild type and muscleless limbs.

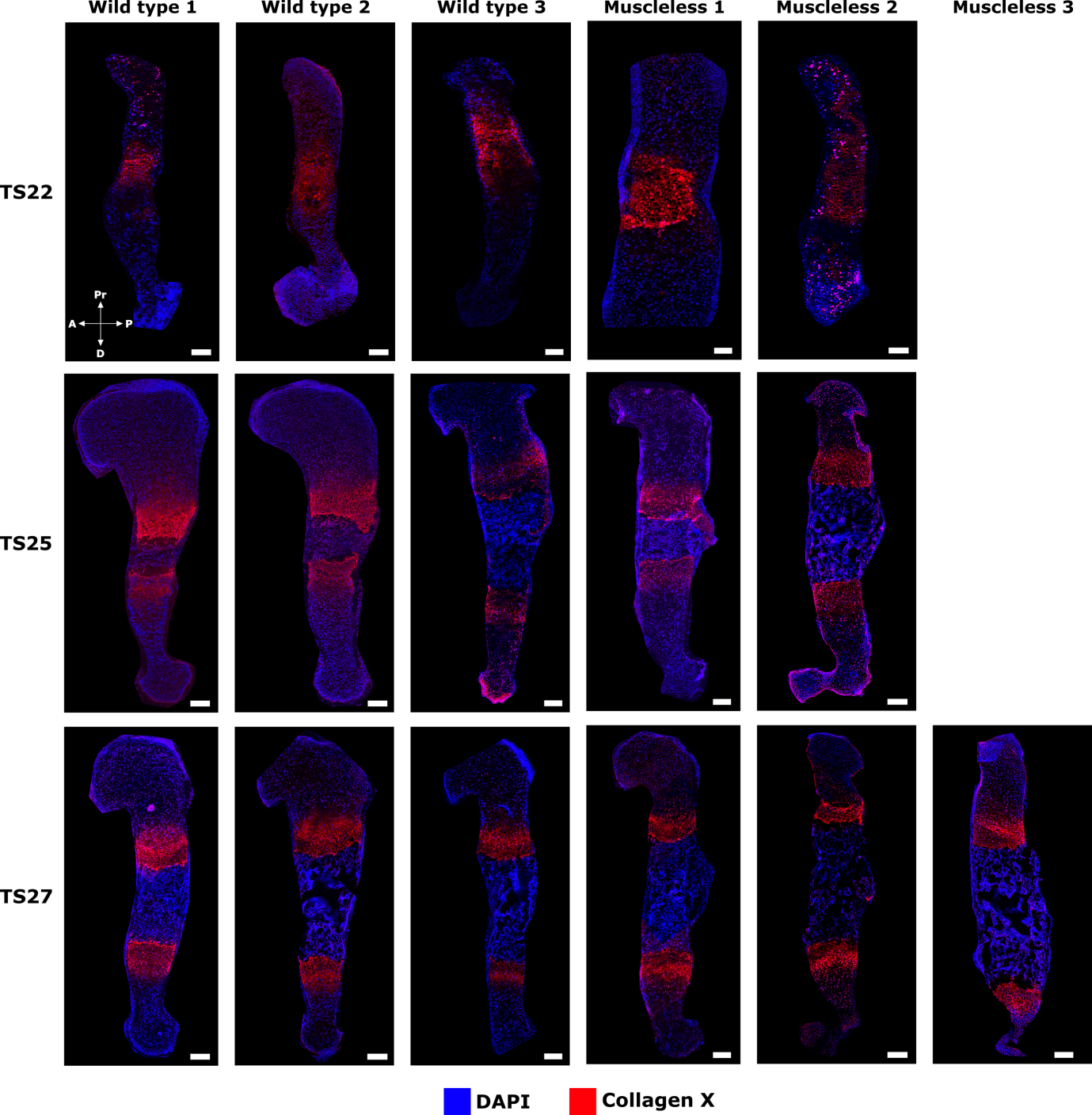

**Figure 7:** Immunofluorescence images of TS22, TS25 and TS27 wild type and muscleless limb humerus stained with collagen X antibody. Scale bars represent 200 μm.

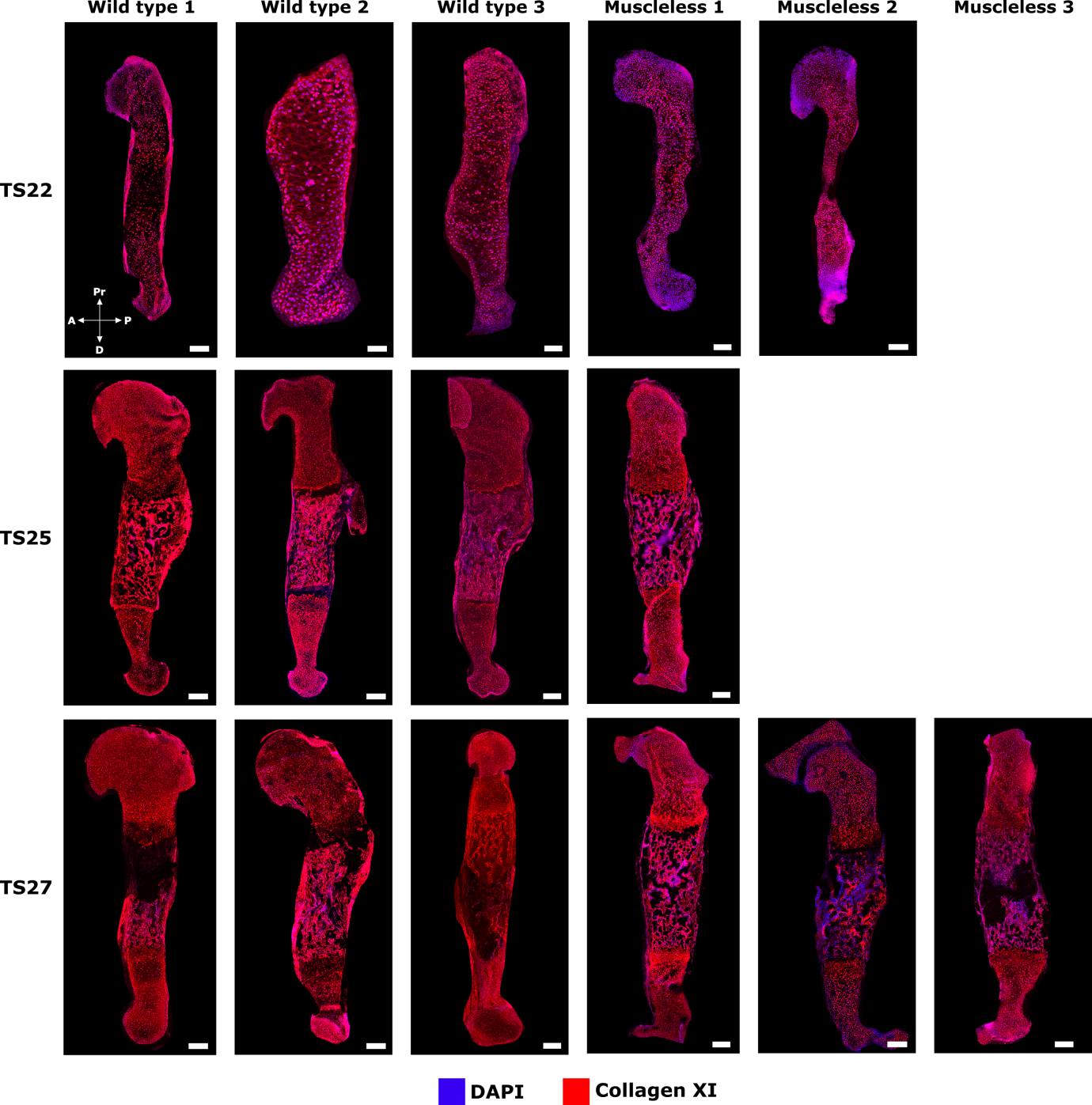

**Figure 8:** Immunofluorescence images of TS22, TS25 and TS27 wild type and muscleless limb humerus stained with collagen XI antibody. Scale bars represent 200 μm.

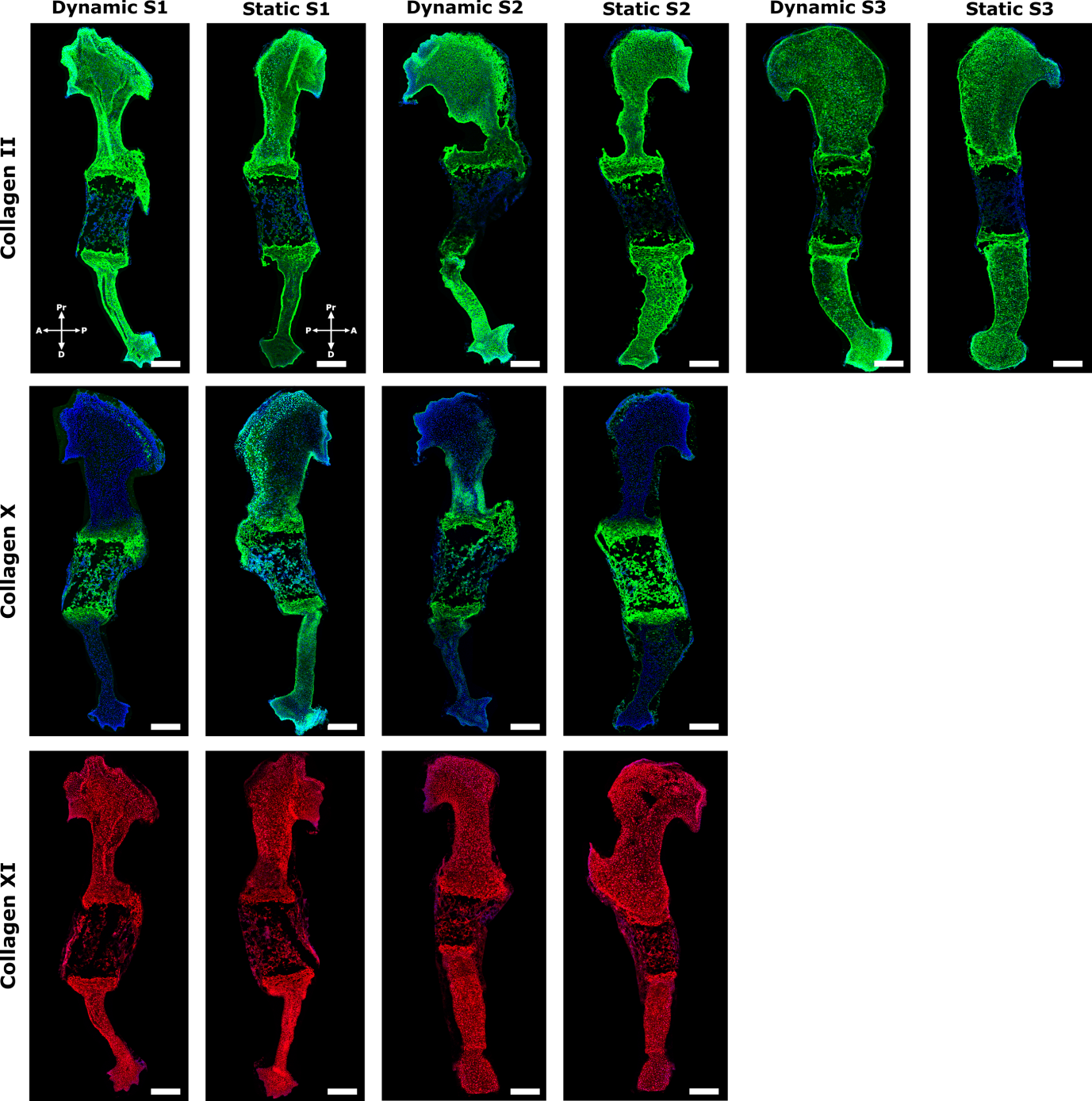

**Figure 9:** Immunofluorescence images of e15.5 wild type humerus cultured under dynamic or static conditions and stained with collagen II, X and XI antibodies. The scale bar is 200 µm.

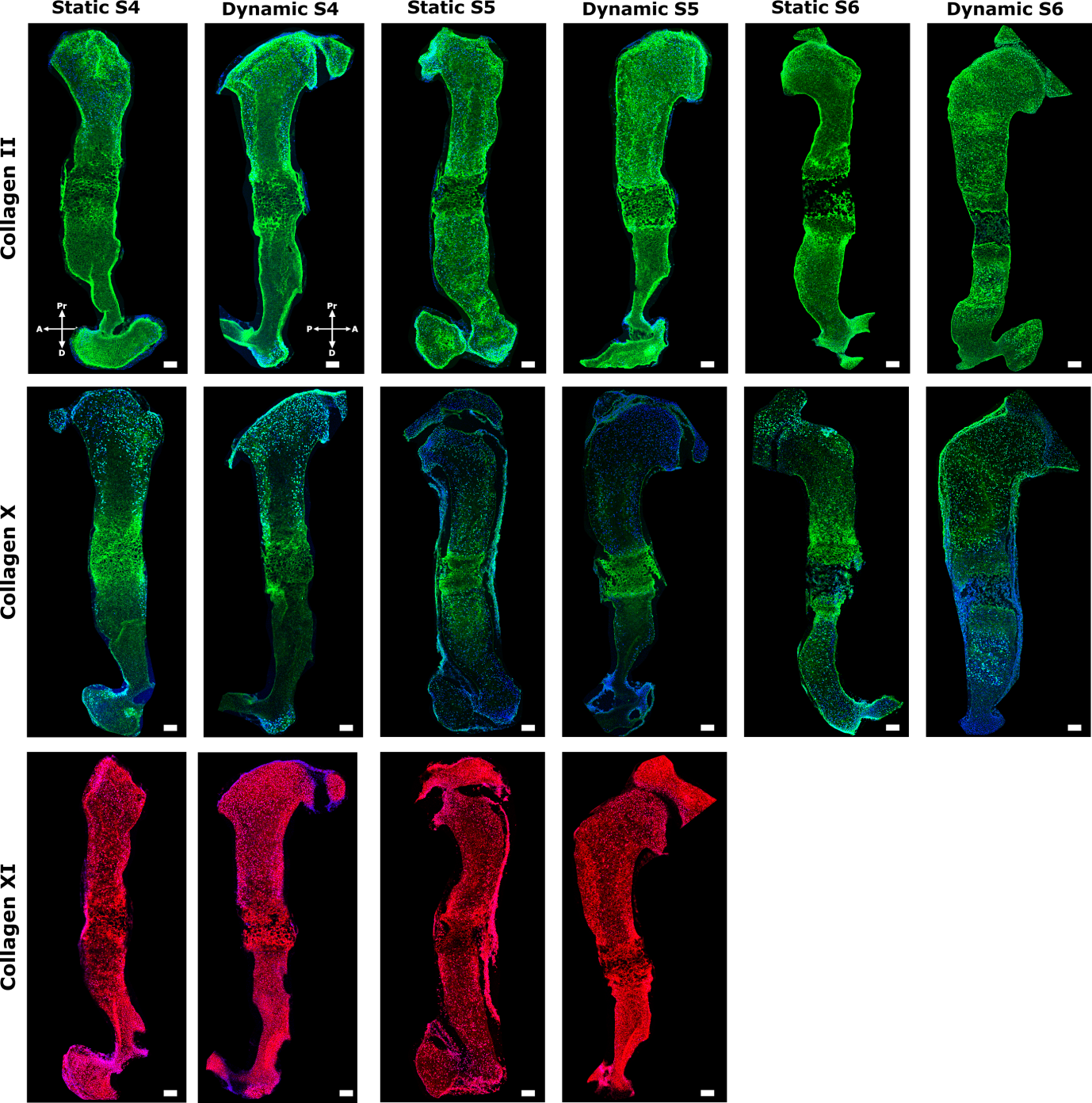

**Figure 10:** Immunofluorescence images of e15.5 musclesless humerus cultured under dynamic or static conditions and stained with collagen II, X and XI antibodies. The scale bar is 100 µm.
